# Autoacetylation-mediated phase separation of TIP60 is critical for its functions

**DOI:** 10.1101/2023.05.06.539700

**Authors:** Shraddha Dubey, Himanshu Gupta, Ashish Gupta

**Affiliations:** Centre of Excellence in Epigenetics, Department of Life Sciences, Shiv Nadar Institution of Eminence, Deemed to be University, Delhi-NCR

**Keywords:** TIP60, autoacetylation, oligomerization, intrinsically disordered region, liquid-liquid phase separation

## Abstract

TIP60 is an important lysine acetyl transferase protein that participates in various essential cellular activities by catalyzing the post-translational acetylation of lysine residues on histones and various non-histone protein substrates. TIP60 typically localizes to the nucleus in a punctate foci pattern, although defining factors and mechanisms regulating the assembly of TIP60 foci and their spatial distribution inside the nucleus are not understood. In the present study, we report that TIP60 can undergo phase separation to form liquid like droplets in the nuclear compartment, which is facilitated by the presence of an intrinsically disordered region (IDR) located between its chromodomain and catalytic domain. Importantly, we identified that autoacetylation on lysine 187, located within the IDR region of TIP60, is important for nuclear localization, oligomer formation and phase separation. Finally, we observed that the phase separation of TIP60 promotes its interaction with its partner proteins and actively contribute to its cellular functions.

## INTRODUCTION

Proteins are the most diverse and abundant of all the organic macromolecules existing in the cell, and perform a variety of functions. The localization of proteins inside the cells never remain static. To function normally and respond adequately to the existing cellular conditions, cells regulate the activities of various proteins by selectively locating them to discrete subcellular compartments. Intracellular compartmentalization of cellular components such as cytoplasm, nucleus, mitochondria, lysosomes, Golgi bodies and endoplasmic reticulum as membrane-enclosed organelles facilitates the creation of specific microenvironments comprising of distinct sets of proteins and other macromolecules. This not only enables the cells to perform distinct cellular functions and carry out complex multistep biochemical reactions where they are required to take place within the cell in the given cellular context, in a spatially and temporally controlled manner. Similarly, confinement of the genetic material and several nuclear factors in the nucleus is crucial to maintain the integrity of genetic material and regulation of gene expression. Unlike other membrane-bound organelles, nuclear compartment may contain a variety of membraneless macromolecular structures such as nucleoli, Cajal bodies, nuclear speckles, paraspeckles, and promyelocytic leukemia protein (PML) bodies which are typically referred to as nuclear bodies^1,2^. Formation of distinct nuclear bodies allows the cells to spatially and temporally segregate a range of nuclear activities such as rRNA biogenesis, small nuclear RNPs, gene regulation, mRNA splicing, nuclear pore complex trafficking and DNA damage repair by inducing heterogeneous positioning of nuclear proteins either alone or along with the nucleic acids, in a context-dependent manner. Several studies have emphasized that spatiotemporal arrangement or positioning of chromosomes to a subnuclear territory allows their dynamic association with varied kind of nuclear proteins, RNA or lncRNAs and thus contributes to the regulation of gene expression^3–5^.

The assembly of non-membrane bound biomolecular condensates is governed via a process called as liquid-liquid phase separation (LLPS) of specific types of proteins known as scaffolds^6,7^. Depending on their composition, these can have solid, gel or liquid-like properties. These structures can be highly dynamic and reversible in nature and can rapidly assemble and disassemble. The ability of cells to form biomolecular condensates by phase separation comes with lot of physiological advantages for the cells. Despite the absence of a membrane or lipid boundary these transient and non-rigid structures maintained as separate and distinct entities within the nucleoplasm allow for dynamic exchange as well as low-specificity interactions of components within the nucleoplasm, at the same time prevent unfavourable interactions^1,3,4^. Nuclear speckle (NS) is one such non-membranous nuclear structure assembled by LLPS-based mechanisms which mainly consist of RNA-binding proteins, splicing factors and lncRNAs and is considered to be hub of active gene transcription although exact mechanism how they regulate gene transcription is still elusive^8,9^. Thus understanding the relationship among nuclear bodies and their constituent macromolecules and their association with three-dimensional organization of genome is important to comprehend the gene expression regulation mechanism to understand how cell responds to the ever changing altering environment.

Until recently, it was believed that proteins use their structured domains to perform their functions, although recent studies have now begun to understand and explore the importance of unstructured regions present in many proteins that typically do not form stable three-dimensional structure of their own^10–12^. These unstructured domains are called intrinsically disordered regions (IDR) and these proteins are called intrinsically disordered proteins (IDP). Notably, condensate-forming proteins frequently contain IDRs which are involved in energetically favourable multivalent intermolecular protein-protein interactions required for phase separation and play important role in regulating enzyme reactions and metabolism within generated condensates^13–15^. Several examples of proteins undergoing phase separation are now known such as histone H2A, OCT4, estrogen receptor, yeast transcription factor GCN4, CTD of Pol II, HP1α and its homolog in Drosophila (HP1), nephrin-nck-n-WASP, DCP2 (mRNA-decapping enzyme subunit 2), EDC3 (enhancer of mRNA-decapping protein 3), NPM1 (nucleolar protein nucleophosmin)^16–19^. Although now there are ample examples of proteins having IDR forming LLPS *in vitro* or *in vivo*, the mechanism regulating these LLPS and their functional significance is still obscure.

TIP60 is an essential lysine acetyl transferase protein known to regulate several important cellular functions including maintenance of genomic integrity, particularly due to its well-recognized involvement in the DNA damage repair process^20–22^. In response to DNA damage, TIP60 can acetylate ATM kinase to help activate the repair pathway and acetylate p53 that assist the cell to decide to either transactivate genes for cell cycle checkpoint activation or for apoptosis depending on the extent of damage^20,22^. TIP60 also contribute to maintain genome integrity by ensuring proper segregation of chromosomes during mitosis through controlling centriole duplication^23^. TIP60 is known to interact with many cellular proteins and can equally acetylate both histone as well as non-histone proteins^24^. Transcriptional activities and cellular function of several nuclear receptors like androgen receptor, estrogen receptor are often regulated by TIP60 in a context-dependent manner^24^. Recently, we have demonstrated the involvement of TIP60 along with PXR in promoting wound healing by modulating the actin reorganization and filopodia formation^25^. TIP60 protein comprise of two characteristic domains common to members of MYST family of HAT proteins: an NH_2_-terminal chromodomain and a highly conserved MYST domain accountable for its catalytic activity. In addition, a short nuclear receptor box (NRB) region is also present near the C-terminal end of TIP60 which helps to mediate its association with many nuclear receptors. TIP60 is generally regarded as a nuclear protein that predominantly distributes in discrete nuclear foci pattern however what mediates the formation of TIP60 foci in the nucleus is not known. Since recent studies have demonstrated the role of liquid-liquid phase separation (LLPS) of proteins and nucleic acids or both in the assembly of various membraneless nuclear structures such as the nucleolus, nuclear paraspeckles, as well as Cajal bodies, we wanted to investigate the role of phase separation as a mechanism behind the formation of the TIP60 punctate foci inside the nuclear compartment. In the present work, we have shown that TIP60 undergoes liquid liquid phase separation mediated by IDR region situated between its chromodomain and MYST domain. Our findings suggest that catalytic activity and chromatin binding of TIP60 plays central role in its IDR-mediated phases separation and autoacetylation of TIP60 at lysine 187 is critical for its phase separation in the nucleus. Divulging further to understand the importance of phase separation for TIP60, we revealed that TIP60 can interact with its interacting partners only in phase separated conditions and investigation of cancer-associated mutations revealed that TIP60’s phase separation is critical for its functions.

## RESULTS

### TIP60 undergo liquid-liquid phase separation mediated through its intrinsically disordered region

TIP60 is known to make punctate foci in the nucleus and we were interested to examine whether these punctate nuclear foci of TIP60 were actually formed by liquid-liquid phase separation (LLPS). For this, we transfected Cos-1 cells with RFP-TIP60 (Wild-type) and subsequently treated these cells with 1, 6-Hexanediol (aliphatic alcohol, which is known to interfere with weak hydrophobic interactions and lead to dissolution of LLPS assemblies both *in vitro* and in the cell) followed by live cell imaging. Results showed significant disruption of TIP60’s foci inside the nucleus by 1, 6-Hexanediol (**Figure 1A**). Fluorescence recovery after photobleaching (FRAP) is used as a standard technique to determine the mobility and dynamic property of these phases separated droplets, whether liquid or solid in nature. Cos-1 cell were transfected with RFP-TIP60 and TIP60 nuclear puncta were photobleached followed by measurement of fluorescence signal. Result showed recovery of fluorescence signal post bleaching with average showing that TIP60 droplets inside the nucleus are dynamic in nature (**Figure 1B**). Together, these findings infer that TIP60 might undergo LLPS-mediated condensate formation.

**Figure 1.**
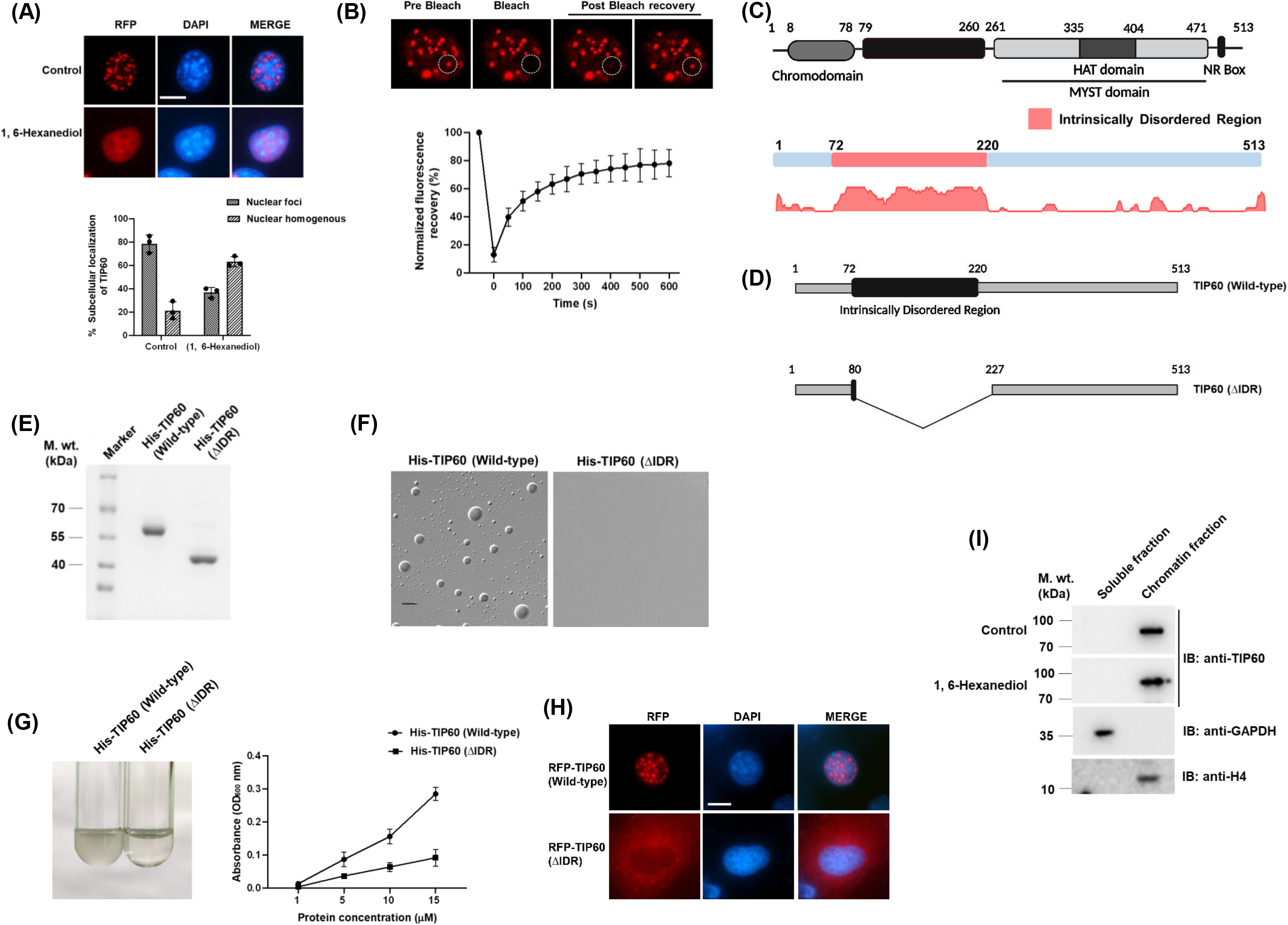
TIP60 undergoes liquid-liquid phase separation through its IDR. **(A)** Presence of 1, 6-Hexanediol disrupts TIP60 nuclear foci. Cos-1 cells transfected with RFP-TIP60 (Wild-type) were treated with either 3.5% 1, 6-Hexanediol or control (water) followed by live cell imaging. RFP panel shows expression of RFP-tagged TIP60. DAPI was used to visualise the nucleus. Scale bar represents 10 µm. Graph (from three independent experimental replicates with SD) generated using GraphPad Prism 8 software, represents percentage of cells showing intranuclear organization of RFP tagged TIP60 in presence or absence of 1, 6-Hexanediol. **(B)** TIP60 nuclear puncta are dynamic liquid droplets. Cos-1 cells were transfected with RFP-TIP60 (Wild-type) and FRAP analysis of live cells were performed. The selected region of interest (ROI) was bleached with 561 nm laser at 100% laser power for 4 seconds followed by post-bleach recovery for 10 minutes. Fluorescence images are of photobleaching experiment and area bleached is depicted by dotted circle. Graph depicts normalized fluorescence recovery (%) for 20 independent puncta’s with SD. Graph is generated using GraphPad Prism 8 software. **(C)** Schematic representation of full-length TIP60 (upper panel) with its different domains: Chromodomain (8-78 amino acids), MYST domain (261-471 amino acids) and NR box (489-493 amino acids). IDR (72-220 amino acids) identified using MobiDB software within TIP60 is highlighted in pink colour. Diagram is created with BioRender.com. (**D)** Schematic diagram showing IDR deleted TIP60. IDR region of TIP60 (from 81 to 226 amino acids) was deleted by overlapping PCR method to generate pET28a-TIP60 (IDR) construct. Diagram is created with BioRender.com. **(E)** Coomassie gel image showing His-TIP60 (Wild-type) and His-TIP60 (ΔIDR) proteins purified from *E. coli*. **(F)** Differential interference contrast (DIC) images representing phase separated liquid droplets of His-TIP60 (Wild-type) and His-TIP60 (ΔIDR) with PEG 8000 as a crowding agent *in vitro*. Scale bar represent 5 µm. **(G)** Representative image showing the variation in turbidity of His-TIP60 (Wild-type) and His-TIP60 (ΔIDR) samples at 15 µM protein concentration in presence of 15% PEG 8000. Graph represents average value of three independent experiments (with SD) for the turbidity assay performed with purified recombinant His-TIP60 (Wild-type) and His-TIP60 (ΔIDR) proteins at different concentrations (1 µM, 5 µM, 10 µM, 15 µM) with 15% PEG 8000. Absorbance of samples was measured at OD_600_ nm. **(H)** IDR deleted TIP60 fail to form nuclear foci. Cos-1 cells were transfected with RFP-TIP60 (Wild-type) or RFP-TIP60 (ΔIDR) plasmids, followed by their live cell imaging to visualize the formation of nuclear foci. Blue colour depicts nucleus stained with DAPI while red colour shows the intracellular expression of RFP tagged TIP60. Scale bar is 10 µm. **(I)** 1, 6-Hexanediol does not affect chromatin binding ability of TIP60. Cos-1 cells transfected with RFP-TIP60 (Wild-type) were subjected to 3.5% 1, 6-Hexanediol treatment followed by subcellular fractionation. Isolated fractions were proceeded for Western blotting using anti-TIP60, anti-GAPDH and anti-H4 antibodies.

TIP60 is composed of different domains including an N-terminal chromodomain, a centrally located MYST domain and a nuclear receptor box (NRB) region located at C-terminal end (**Figure 1C**). To predict the ability of TIP60 to form LLPS, we computationally analyzed its sequence using MobiDB^26–28^ software to look for the presence of intrinsically disordered region (IDR) in TIP60. Interestingly, we identified the presence of an IDR located between amino acid residues 72 to 220, flanked on one side by N-terminal chromodomain and by MYST domain on another side (**Figure 1C**). We also used other software such as dSCOPE^29^ and PONDR to validate the presence of IDR in TIP60 and found TIP60 to harbour IDR in the similar region (**Figure S1A and S1B**). To further estimate the ability of TIP60 to form liquid droplets, we used the FuzDrop^30^ software to identify droplet-promoting regions (DPRs) in the TIP60 protein and interestingly, DPR was found to be present between amino acids residues 69 to 219, which completely overlaps with the IDR region of TIP60 (**Figure S1C**).

Once the putative IDR region in TIP60 was identified, we wanted to investigate whether TIP60 can form LLPS and whether the IDR has a role in this. For this, we first purified recombinant TIP60 full-length protein (His-TIP60 (Wild-type)) and IDR-deleted TIP60 protein (His-TIP60 (ΔIDR)) from *E. coli* using affinity chromatography (**Figure 1D & 1E**). To examine phase separation property of TIP60 protein by LLPS under *in vitro* conditions, we treated purified His-TIP60 (Wild-type) protein with different concentrations of PEG 8000 as crowding reagent and examined the formation of liquid droplets by differential interference contrast (DIC) microscopy. Result showed propensity of LLPS-mediated droplet formation by His-TIP60 (Wild-type) protein under examined conditions, which significantly increased in presence of 15% PEG 8000 (**Figure S2A**). To explore the role of TIP60’s IDR in its phase separation, we then examined phase-separated droplet forming ability of His-TIP60 (Wild-type) and His-TIP60 (ΔIDR) proteins with 15% PEG 8000, and found that His-TIP60 (ΔIDR) protein failed to form LLPS-mediated droplets under these conditions (**Figure 1F**).

Since conversion of a protein from soluble state to phase separated state may perturb the turbidity of solution thus measuring variations in turbidity of solution (Turbidity assay) is a biochemical analytical method to study phase separation property of the protein. Thus, we also measured the turbidity (OD_600_) of the solution containing varied concentrations of His-TIP60 (Wild-type) or His-TIP60 (ΔIDR) proteins and result showed significant difference in the turbidity between both the solutions with His-TIP60 (Wild-type) displaying significantly high absorbance compared to His-TIP60 (ΔIDR) (**Figure 1G**). Together these results demonstrate that TIP60 protein undergoes LLPS under *in vitro* condition, mediated by its IDR.

Next we wanted to examine the role of IDR on TIP60’s nuclear foci formation by live cell imaging experiments. For this, we cloned TIP60 (ΔIDR) ORF into pDsRed vector and confirmed its expression by Western blot analysis (**Figure S2B**). Cos-1 cells were transfected with RFP-TIP60 (Wild-type) or RFP-TIP60 (ΔIDR) plasmids followed by live cell imaging to monitor their intracellular localization pattern. Remarkably, we observed that unlike RFP-TIP60 (Wild-type) protein that formed punctate foci inside the nucleus, RFP-TIP60 (ΔIDR) protein did not form any punctate foci and got accumulated in the cytosol (**Figure 1H**). These results showed role of IDR in TIP60’s ability to form LLPS-mediated nuclear foci.

We also wanted to ensure that 1, 6-Hexanediol-mediated disruption of TIP60’s foci inside the nucleus is due to dissolution of its LLPS and not due to variation in TIP60 protein levels following 1, 6-Hexanediol treatment, we performed Western blot analysis with same set of samples and found no observable change in TIP60 protein levels under these conditions (**Figure S2C**). Since TIP60 is known to remain chromatin bound^25^, so we wanted to determine the impact of 1, 6-Hexanediol-mediated nuclear foci disruption of TIP60 on its chromatin binding ability. For this, we performed subcellular fractionation assay with Cos-1 cells transfected with RFP-TIP60 and treated with 1, 6-Hexanediol. Western blot result detected the presence of TIP60 exclusively in chromatin-bound fraction both in control and in 1, 6-Hexanediol treated samples indicating that TIP60 LLPS are formed in chromatin-bound state (**Figure 1I**).

### Autoacetylation of lysine 187 situated within IDR is essential for TIP60’s nuclear import and LLPS

Since our results show that TIP60’s nuclear foci are formed due to phase separation and in a previous study from our laboratory, we showed that HAT mutant of TIP60 remains homogeneously distributed inside the nucleus^31^. Therefore, to examine if TIP60’s catalytic activity is important for its nuclear foci formation, we performed live cell imaging experiments to examine the localization of TIP60 in presence of NU9056 (inhibitor of TIP60’s catalytic activity). For this, Cos-1 cells were transfected with RFP-TIP60 (Wild-type) and the transfected cells were treated either with DMSO (vehicle control) or NU9056. Imaging results showed that inhibiting TIP60’s catalytic activity alters its nuclear organization and drives it into a homogeneously distributed pattern (**Figure 2A**). Having observed that the catalytic activity of TIP60 is important for its foci formation and knowing that phase separation of TIP60 occurs in the chromatin-bound state (**Figure 1I**), we set out to investigate the relationship between the catalytic activity of TIP60 and its chromatin binding ability with respect to its phase separation. For this, Cos-1 cells were transfected with RFP-TIP60 (Wild-type) or RFP-TIP60 (HAT mutant) and soluble and chromatin bound fractions were prepared followed by Western blot analysis. The results showed that in comparison to RFP-TIP60 (Wild-type) which exclusively remained in chromatin fraction, RFP-TIP60 (HAT mutant) drastically lost its chromatin binding ability as significant proportion of this mutant protein was detected in the soluble fraction (**Figure 2B**).

**Figure 2.**
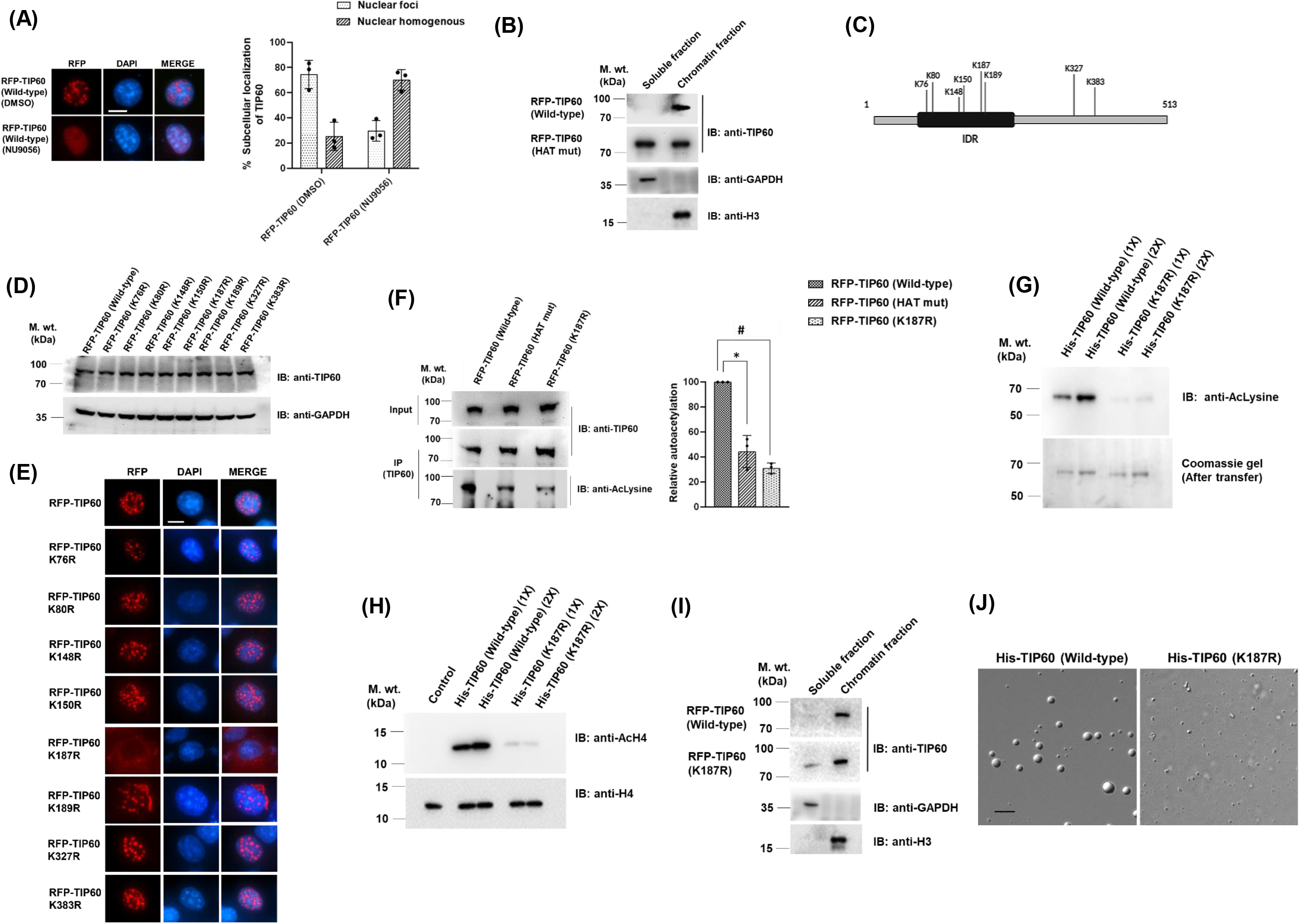
Autoacetylation of lysine 187 situated within IDR of TIP60 regulate its nuclear localization and phase separation. **(A)** Inhibition of TIP60’s catalytic activity by NU9056 activity disrupts its nuclear foci. Cos-1 cells transfected with RFP-TIP60 plasmid were treated with NU9056 or DMSO (control) followed by live cell imaging. Red colour depicts expression of RFP tagged protein and blue colour depicts nucleus stained with DAPI. Scale bar represents 10 µm. Graph shows percentage of RFP-TIP60 intranuclear organization in transfected cells in presence of NU9056 or DMSO from the data obtained from three independent experimental replicates with SD. **(B)** TIP60’s catalytic activity is important for its chromatin binding. Cos-1 cells transfected with RFP-TIP60 (Wild-type) or RFP-TIP60 (HAT mutant) were subjected to subcellular fractionation. Western blot analysis was performed with isolated fractions with anti-TIP60, anti-GAPDH and anti-H3 antibodies. **(C)** Schematic diagram depicting autoacetylation lysine residues sites in TIP60 protein. Residue numbers are given according to isoform 2 of TIP60. Diagram is created using BioRender.com. **(D)** Western blot image showing the expression of generated TIP60 autoacetylation mutant constructs. Cos-1 cells were transfected with RFP-TIP60 (Wild-type) or various autoacetylation mutant plasmids. Harvested cell lysates were resolved on SDS-PAGE followed by Western blot analysis using anti-TIP60 or anti-GAPDH antibody. **(E)** Autoacetylation of lysine 187 located within IDR is essential for nuclear localization of TIP60. RFP-TIP60 (Wild-type) or its autoacetylation mutant plasmids were transfected in Cos-1 cells followed by live cell imaging. Blue colour represents nucleus stained with DAPI and red colour shows expression of RFP tagged TIP60 proteins. Scale bar represents 10 µm. **(F)** TIP60 (K187R) shows significantly reduced autoacetylation activity. Immunoprecipitation of TIP60 protein was performed with the lysates of Cos-1 cells transfected with indicated plasmids followed by Western blot analysis using anti-acetylated lysine or anti-TIP60 antibody. Graph (constructed using GraphPad Prism 8 software) represents the autoacetylation level of TIP60 proteins for the quantitated values obtained from three independent experimental replicates (SD), taking autoacetylation of TIP60 (Wild-type) level as 100%. p value for RFP-TIP60 (Wild-type) vs RFP-TIP60 (HAT mutant) is 0.0171 (*) and for RFP-TIP60 (Wild-type) vs RFP-TIP60 (K187R) is 0.0011 (#). **(G)** His-TIP60 (K187R) exhibits significantly decreased autoacetylation levels. *In vitro* autoacetylation assay was performed using recombinant purified His-TIP60 (Wild-type) or His-TIP60 (K187R) proteins. Samples were processed for Western blot analysis using anti-acetylated lysine antibody. Coomassie gel represents the loading of reaction samples. **(H)** TIP60 (K187R) mutant shows diminished histone H4 acetylation activity. His-TIP60 (Wild-type) and His-TIP60 (K187R) proteins were incubated with histone H4 peptides and samples were resolved in 15% SDS-PAGE followed by Western blotting using anti-acetylated H4 and anti-H4 antibody. **(I)** TIP60 (K187R) mutant retains its chromatin binding property. Cos-1 cells transfected with RFP-TIP60 (Wild-type) or RFP-TIP60 (K187R) were subjected to subcellular fractionation and isolated fractions were subsequently proceeded for Western blotting using anti-TIP60, anti-GAPDH and anti-H3 antibody. **(J)** DIC images showing i*n vitro* liquid-liquid phase separation of His-TIP60 (Wild-type) and His-TIP60 (K187R) proteins facilitated with PEG 8000 as a crowding agent. Scale bar is equivalent to 5 µm.

Since, TIP60 is known to autoacetylate itself which is dependent on its catalytic activity^32,33^, we wanted to determine the impact of TIP60’s autoacetylation on its LLPS-mediated nuclear foci formation. For this, we selected autoacetylation sites which are found to be conserved in all the isoforms of TIP60 to rule out any isoform specific predisposition. Eight such autoacetylation sites were identified in TIP60 and acetylation null point mutant were generated for each of these sites by converting lysine into arginine (**Figure 2C**). Western blot analysis was performed to examine the expression of generated autoacetylation mutants of TIP60 cloned into pDsRed vector and result showed that all the generated mutants expressed full-length protein at their expected size (**Figure 2D**). Further, to determine the impact of these mutations on TIP60’s intracellular localization, we transfected Cos-1 cells with these autoacetylation mutant plasmids and performed live cell imaging. Results showed that except RFP-TIP60 (K187R) mutant which showed completely cytosolic localization similar to TIP60 (ΔIDR) protein, all other autoacetylation mutants behaved similar to wild type TIP60 protein and could formed punctate nuclear foci (**Figure 2E**).

On observing the impact of lysine 187 mutation on TIP60’s phase separation *in vivo*, we wanted to further examine the impact of this mutation on its autoacetylation and histone substrate acetylation. Since our results showed that phase separation of TIP60 requires its efficient loading onto the chromatin under *in vivo* conditions we also wanted to examine the status of chromatin binding ability of this mutant. To examine the impact on lysine 187 mutation on TIP60’s autoacetylation and for its comparison with wild-type and HAT mutant of TIP60 protein, we performed autoacetylation assay using Cos-1 cells transfected with RFP-TIP60 (Wild-type) or RFP-TIP60 (K187R) mutant or RFP-TIP60 (HAT mutant). TIP60 protein was immunoprecipitated using anti-TIP60 antibody followed by Western blot analysis with anti-TIP60 antibody or anti-acetylated lysine antibody. Results showed significant decrease in acetylation level of TIP60 HAT mutant and TIP60 (K187R) mutant proteins (**Figure 2F**). We performed similar assay with recombinant His-TIP60 (Wild-type) and His-TIP60 (K187R) protein purified from bacteria and drastic decrease in auto-acetylation level of His-TIP60 (K187R) mutant protein was observed by Western blot analysis (**Figure 2G**). Next, to examine whether lysine 187 mutation had any impact on TIP60’s catalytic activity for histone substrate we performed HAT assay with purified recombinant His-TIP60 (Wild-type) or TIP60 (K187R) mutant proteins using histone H4 peptide as substrate, followed by their Western blot analysis with anti-acetylated H4 antibody. The result showed that lysine (K187R) mutant significantly lost its histone acetylation ability suggesting that lysine 187 has considerable impact not only on its autoacetylation but also affected TIP60’s catalytic activity for histone substrate under *in vitro* conditions (**Figure 2H**). To determine the impact of lysine 187 mutation on TIP60’s chromatin binding ability, subcellular fractionation assay was performed using Cos-1 cells transfected with RFP-TIP60 (Wild-type) or RFP-TIP60 (K187R) or RFP-TIP60 (HAT mutant) plasmids and subsequently Western blot analysis was performed. To our surprise, the chromatin binding ability of TIP60 (K187R) mutant was found to be intact despite its inability to enter the nucleus, suggesting that autoacetylation of lysine 187 does not have any role to play in chromatin binding of TIP60 protein (**Figure 2I**).

Next, we wanted to determine the status of phase separation for the autoacetylation-defective mutant of TIP60 (K187R), which had failed to enter the nucleus within the cell. The *in vitro* LLPS assay was performed with purified recombinant His-TIP60 (Wild-type) and His-TIP60 (K187R) proteins and the DIC microscopy results showed that compared with TIP60 (Wild-type), we observed a significant reduction in the size of LLPS-mediated droplets of His-TIP60 (K187R), which highlight the importance of the lysine 187 site located within the IDR in regulating the phase separation of TIP60 (**Figure 2J**). Furthermore, we performed Clustal Omega analysis for TIP60 protein sequences from different species and found lysine 187 to be conserved specifically in all the examined vertebrate TIP60 proteins (**Figure S2D**). Together, all these results show that autoacetylation of lysine 187 (located within IDR) act as a regulator of TIP60’s phase separation.

### Autoacetylation of lysine 187 plays important role in TIP60 oligomerization

Having determined the role of lysine 187 autoacetylation in TIP60’s phase separation, we wanted to explore the mechanism by which lysine 187 may facilitate TIP60 LLPS. As it has been shown that oligomerization can favour the ability of proteins to form LLPS^34^, we wanted to determine whether the TIP60 protein can form high molecular weight oligomers and does lysine 187 has any role in this. To examine and compare the oligomer forming property of TIP60 and its mutant proteins, we performed glutaraldehyde cross linking assay using His-TIP60 (Wild-type), His-TIP60 (K187R) or His-TIP60 (HAT mutant) purified recombinant proteins (**Figure S3A**). Western blot analysis was performed and the results showed that the TIP60 (Wild-type) protein can form high molecular weight oligomers, mainly trimers (**Figure 3A**). In addition to trimers, minor populations of TIP60 dimers and other high molecular weight oligomers were also observed. Notably, under similar experimental conditions, we observed a drastically reduced oligomerization property for the TIP60 (K187R) and TIP60 (HAT mutant) proteins, suggesting that lysine 187 autoacetylation is important for TIP60’s oligomer formation (**Figure 3A**). Further we performed *in vitro* interaction studies to examine whether the loss of oligomerization ability of TIP60 (K187R) mutant protein, affects interaction between TIP60-TIP60 protein. For this *E. coli* BL21 DE3 strain was cotransformed with pGEX6p2-TIP60 (Wild-type) and pET28a-TIP60 (Wild-type) or pET28a-TIP60 (K187R) plasmids. GST-TIP60 (Wild-type) protein was purified under native conditions using gluthathione S-sepharose beads in a bead-bound state. Subsequently, Western blot analysis was performed with purified bead-bound protein samples using anti-His antibody which showed copurification of His-TIP60 (Wild-type) protein along with GST-tagged TIP60 (Wild-type) protein, however under similar conditions, we found no significant copurification of His-TIP60 (K187R) protein (**Figure 3B).** This revealed that lysine 187 autoacetylation is important for TIP60–TIP60 protein interaction.

**Figure 3.**
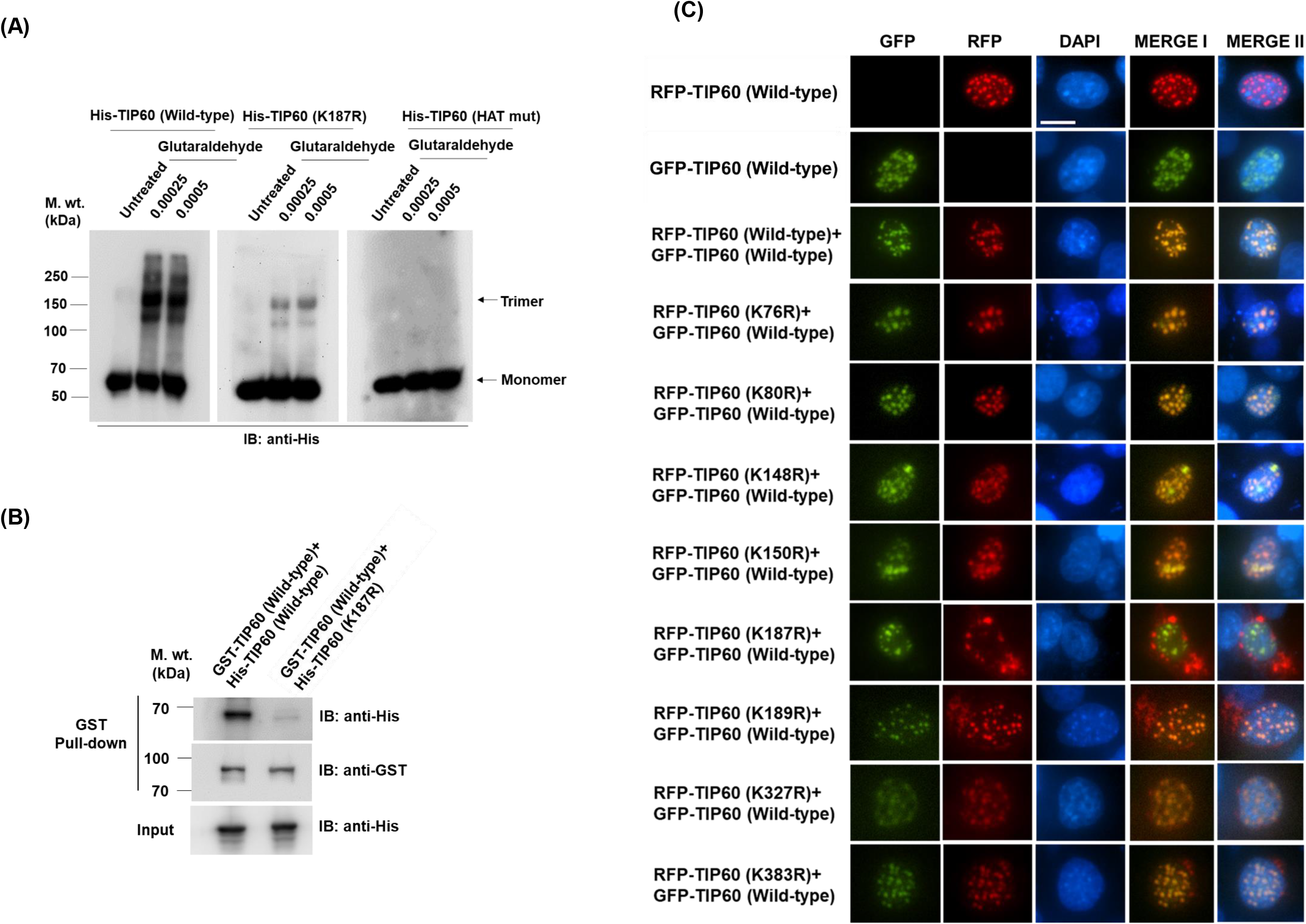
TIP60 autoacetylation at lysine 187 residue is essential for its oligomerization. **(A)** TIP60 predominantly forms trimer. Recombinant purified His-TIP60 (Wild-type), His-TIP60 (K187R) and His-TIP60 (HAT mutant) protein samples incubated with glutaraldehyde followed by resolution of reaction samples in 8% SDS-PAGE. Western blotting was performed using anti-His antibody. **(B)** TIP60’s 187 lysine residue is critically important for TIP60-TIP60 interaction. *E. coli* BL21 DE3 cells were transformed with indicated plasmids followed by expression of recombinant proteins. GST-TIP60 protein was purified using glutathione S-transferase beads in bead bound form and samples were processed for Western blot analysis using anti-GST and anti-His antibody. **(C)** RFP-TIP60 (K187R) mutant failed to colocalize with TIP60 (Wild-type) *in vivo*. Live cell imaging was performed to visualise the localization of GFP-TIP60 (Wild-type) and RFP-TIP60 (Wild-type or autoacetylation mutant) plasmids transfected in Cos-1 cells. DAPI was used to stain the nucleus. Red colour shows the expression of RFP tagged TIP60 (Wild-type or autoacetylation mutants) proteins while green colour shows the localization pattern of GFP tagged TIP60 protein. Merge I represents the colocalized images of GFP and RFP panel while merge II shows the GFP, RFP and DAPI colocalization. Scale bar is equivalent to 10 µm.

Since we have already shown in this study that TIP60 (K187R) mutant fails to translocate into the nucleus and subsequent results showed the importance of the same lysine residues in TIP60-TIP60 protein interaction and its oligomerization, we further wanted to investigate the effect of TIP60 oligomer formation on its phase-separated nuclear foci formation. Through live cell imaging experiment, we examined the intracellular localization and foci formation of TIP60 (K187R) mutant in presence of its wild-type form in the cell. For this, we first generated TIP60 clone in pEGFP vector and confirmed its expression by Western blotting (**Figure S3B**). To determine its intracellular localization, live cell imaging was performed for Cos-1 cells transfected with GFP-TIP60 (Wild-type) clone and result showed that GFP-tagged TIP60 formed nuclear punctate foci in a similar manner to the RFP-tagged TIP60 in the nucleus (**Figure 3C**).

Next, we cotransfected Cos-1 cells with RFP-tagged TIP60 (Wild-type) or its autoacetylation mutant plasmids and GFP-tagged TIP60 (Wild-type) plasmids and performed live cell imaging to monitor their localization. We observed clearly visible colocalized foci of RFP-tagged TIP60 with GFP-tagged TIP60 in the nucleus, and similarly, all other autoacetylation mutants except TIP60 (K187R) mutant formed colocalized nuclear foci with GFP tagged TIP60 (**Figure 3C**). However, the presence of GFP-tagged wild-type TIP60 protein could not induce nuclear import of the RFP-TIP60 (K187R) mutant protein, and both these proteins remain distributed individually in the nucleus and cytosol, respectively. These results reaffirm our observation that lysine 187 autoacetylation is critical for TIP60-TIP60 protein interaction and oligomerization, required for its phase separation.

### TIP60 partly assimilate into nuclear speckles

Once the phase separation of TIP60 was confirmed, our next step was to determine the spatial distribution of phase-separated TIP60 protein inside the nucleus. Since TIP60’s foci pattern bears a resemblance with nuclear speckles, we performed immunofluorescence assay in Cos-1 cells to examine the status of TIP60 localization with respect to NS marker SC35. Confocal microscopy result showed partial colocalization between TIP60 and SC35 foci within the nucleus (**Figure 4A**). To further determine the assimilation of TIP60 into NS, we performed immunoprecipitation assay using TIP60 antibody or IgG control from Cos-1 cells followed by Western blot analysis with anti-TIP60 and anti-SC35 antibody. Result showed the presence of SC35 signal in TIP60-immunoprecipitated lysate while no signal of SC35 was detected in IgG immunoprecipitated sample showing presence of TIP60 in the NS (**Figure 4B).**

**Figure 4.**
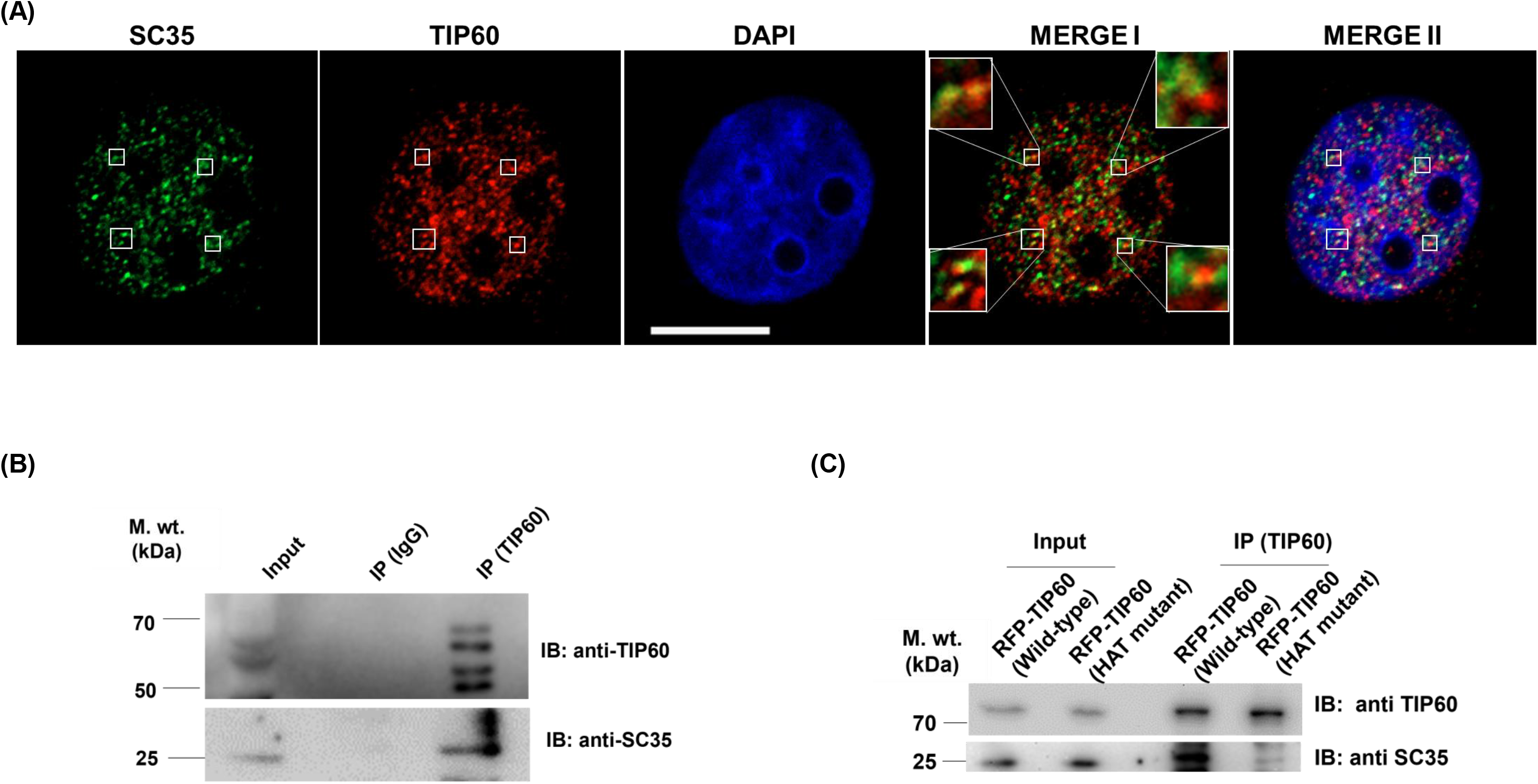
Catalytic activity of TIP60 is essential for its assimilation into nuclear speckles. **(A)** TIP60 partially colocalize with NS marker SC35. Cos-1 cells were subjected to immunofluorescence assay using anti-TIP60 (mice) and anti-SC35 (rabbit) primary antibodies and Alexa fluor 594 anti-mice and Alexa fluor 488 anti-rabbit used as secondary antibodies. Nucleus was stained with DAPI (blue colour). Red colour and green colour depict endogenous localization of TIP60 and SC35 respectively. Enlarged image (inset) show colocalization of TIP60 with SC35. Merge I image show combined signals for TIP60 and SC35 while Merge II depicts combined signals for TIP60, SC35 and DAPI. Scale bar represents 10 µm. **(B)** TIP60 immunoprecipitation pulls down SC35 in Cos-1 cells. Cos-1 cell lysate was subjected to immunoprecipitation using TIP60 antibody or IgG. Bead-bound proteins were resolved in SDS-PAGE and Western blotting was performed with anti-TIP60 or anti-SC35 antibody. **(C)** TIP60’s catalytic activity is important for its interaction with NS. Cos-1 cells were transfected with mentioned plasmids and immunoprecipitation was performed using TIP60 antibody. Western blot analysis was performed for the bead-bound protein samples, using anti-TIP60 and anti-SC35 antibody.

Since TIP60’s catalytic activity was found to be essential for phase separation *in vivo* (**Figure 2A**), to explore whether TIP60’s catalytic activity is also required for its assimilation into NS, we performed immunoprecipitation assay with Cos-1 cells transfected with RFP-TIP60 (Wild-type) or RFP-TIP60 (HAT mutant) using TIP60 antibody. Subsequently, Western blot analysis of the immunoprecipitated samples was performed using anti-TIP60 and anti-SC35 antibody and the result showed that SC35 was successfully pulled down only with TIP60 (Wild-type) immunoprecipitated samples, however, compared to TIP60 (Wild-type) immunoprecipitated samples, very weak signal of SC35 was detected in the TIP60 (HAT mutant) immunoprecipitated samples (**Figure 4C**). Together these findings suggest that TIP60 partially assimilate into NS and its catalytic activity actively contributes in this assimilation.

### TIP60 interacts with its partner protein in its phase separated form

TIP60 is a multifunctional protein known to interact with various cellular proteins to execute its function in a context-dependent manner. Recently, we have shown the role of TIP60 in promoting wound healing process through its association with PXR, thus we chose to examine the role of TIP60’s phase-separation on its interaction with PXR as one of its cellular partners. To test whether TIP60 phase separation has a role in regulating TIP60’s interaction with PXR, we examined & compared the localization and interaction of PXR with Wild-type TIP60 which can phase separate and TIP60 HAT mutant which do not undergo phase separation. For this, we performed live cell imaging experiments using co-transfected Cos-1 cells with GFP-PXR along with RFP-TIP60 (Wild-type) or RFP-TIP60 (HAT mutant) and found that PXR only makes colocalized foci with RFP-TIP60 (Wild-type) however failed to do so with RFP-TIP60 (HAT mutant) (**Figure S4A**). To analyse the interaction between these proteins, we performed immunoprecipitation assay from the cell lysate of these transfected Cos-1 cells using anti-PXR antibody and Western blot analysis of the immunoprecipitated samples showed exclusive interaction of PXR only with RFP-TIP60 (Wild-type) and not with RFP-TIP60 (HAT mutant) (**Figure S4B**). These results suggest that PXR can interact only with phase separated TIP60. Interestingly we also observed that although TIP60 (HAT mutant) when expressed alone remain evenly distributed in the nucleus, however, when co-expressed with TIP60 (Wild-type), it forms colocalized punctate nuclear foci with TIP60 (Wild-type), which indicates that even a single catalytically functional subunit of TIP60 is sufficient to form its oligomer.

Since phase-separated TIP60 has been shown to form oligomers, it was important to investigate whether this phase-separated form of TIP60 maintains its oligomeric form while interacting with its partners or may alter its structure to monomeric form to favour the interaction. To examine these possibilities, we decided to coexpress TIP60 (Wild-type) and TIP60 (HAT mutant) proteins tagged with different fluorescent tags together in combination with PXR in Cos-1 cells. If PXR interacted with oligomeric TIP60 (composed of TIP60 (Wild-type) and TIP60 (HAT mutant) proteins differentially tagged with RFP and GFP), the formation of these colocalized TIP60 foci should have remain intact and no effect would be seen on number of cotransfected cells forming these foci, in presence of PXR. However, if PXR interacted only with monomeric phase-separated TIP60, we would see disruption of colocalized foci (GFP-and RFP-tagged TIP60 (Wild-type) and TIP60 HAT mutants) in the presence of PXR. Live cell imaging of Cos-1 cells transfected with different combinations of plasmids showed no visible disruption of colocalized foci of fluorescence-tagged Wild-type and HAT mutant TIP60 proteins, as shown in Figure 5A, suggesting that PXR interacts exclusively with oligomerized phase-separated TIP60 (**Figure 5A**).

**Figure 5.**
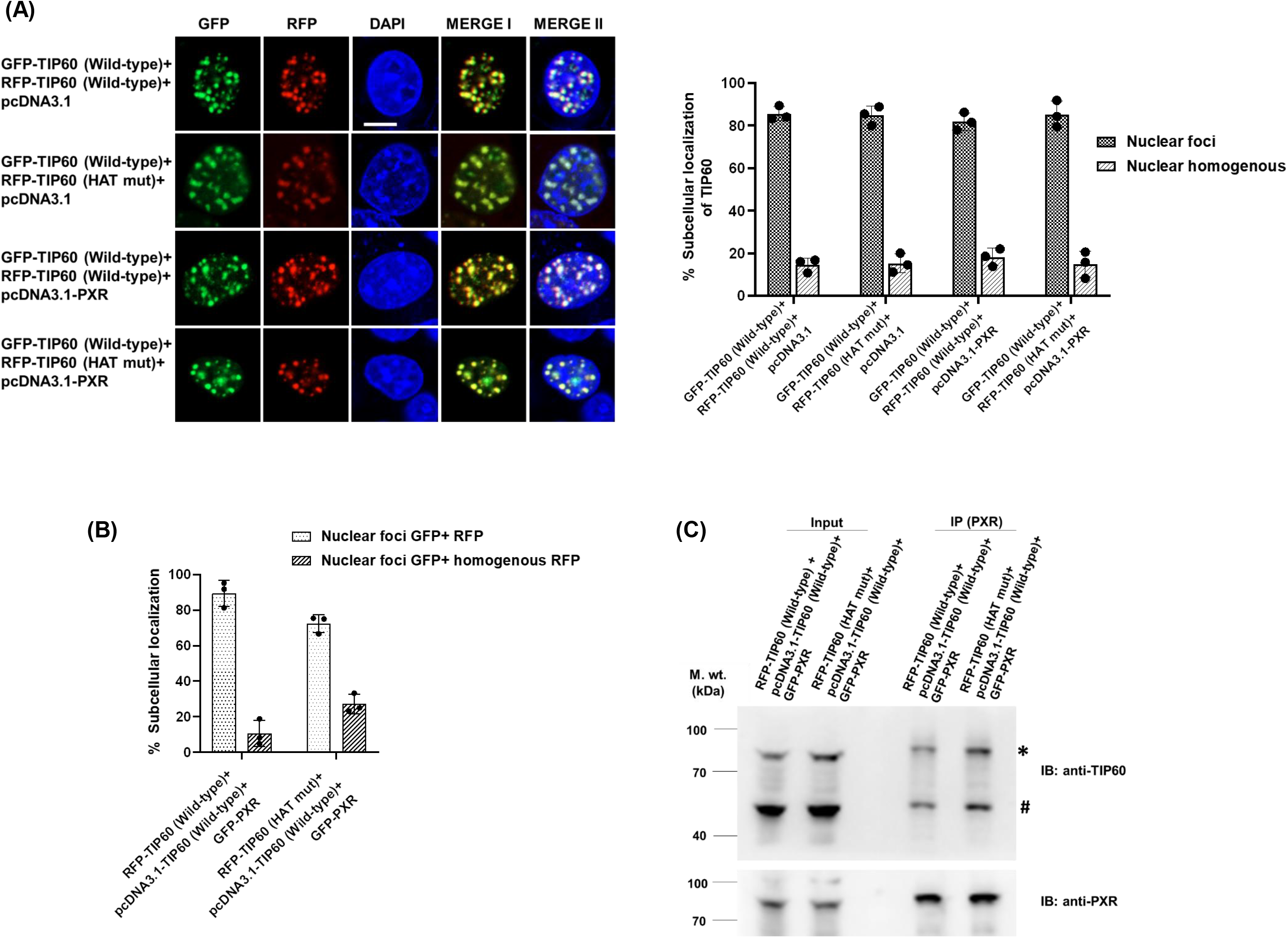
Phase separation of TIP60 is important for its interaction with partner protein. **(A)** PXR interacts with phase-separated TIP60. Cos-1 cells were transfected with the plasmids as mentioned in the figure and live cell imaging was performed. RFP tagged TIP60 (Wild-type) or TIP60 (HAT mutant) are shown in red colour, whereas GFP tagged TIP60 (Wild-type) is shown in green colour. Merge I illustrate combined RFP and GFP signals, while merge II depicts combined RFP, GFP, and DAPI signals. DAPI is used for depicting the nucleus (blue colour). Scale bar is equivalent to 10 µm. Graph depicts percentage localization data of fluorescent tagged proteins obtained from three independent experimental replicates with SD. Graph was generated using GraphPad Prism 8 software. **(B)** Cos-1 cells were transfected with plasmids (as indicated in the figure) followed by live cell imaging for detecting colocalization of fluorescent tagged proteins. Graph was generated using GraphPad Prism 8 software depicting the percentage colocalization of fluorescent-tagged proteins in transfected cells, from three independent experiments with SD. **(C)** PXR interact with phase separated TIP60 oligomers. Cos-1 cells were transfected with the plasmid combinations as depicted in the figure and PXR immunoprecipitation was performed using anti-PXR antibody. Immunoprecipitated samples were resolved in SDS-PAGE followed by Western blotting using anti-TIP60 and anti-PXR antibody. Asterisk (*) shows signal detected for RFP-TIP60 (Wild-type) or RFP-TIP60 (HAT mutant), whereas the hash (#) shows the signal for pcDNA3.1-TIP60 in immunoprecipitated samples.

To rule out any experimental bias, we performed similar experiment again by swapping the tags of the proteins as indicated in Figure 5B. Cos-1 cells were transfected with two different sets of plasmid combinations-RFP-TIP60 (Wild-type), GFP-PXR and pCDNA3.1-TIP60 (Wild-type) or RFP-TIP60 (HAT mutant), GFP-PXR and pCDNA3.1-TIP60 (Wild-type). Live cell imaging results showed that RFP-TIP60 (HAT mutant) and GFP-PXR that could not form foci when co-expressed together, were now able to form colocalized foci in the presence of the Wild-type form of TIP60 which again confirms that PXR can only interact with oligomerized phase-separated TIP60 (**Figure 5B**). Further, to confirm interaction of PXR with oligomerized TIP60, we performed immunoprecipitation assay using anti-PXR antibody from the lysates of Cos-1 cells transfected with two different sets of plasmid combinations-RFP-TIP60 (Wild-type), GFP-PXR and pCDNA3.1-TIP60 (Wild-type) or RFP-TIP60 (HAT mutant), GFP-PXR and pcDNA3.1-TIP60 (Wild-type). Western blot results showed the presence of both TIP60 (Wild-type) and TIP60 (HAT mutant) proteins in PXR-immunoprecipitated samples, conclusively suggesting that PXR interacts only with oligomerized phase-separated TIP60 inside the nucleus (**Figure 5C**).

### TIP60 phase separation is essential to perform its cellular function

Once it is confirmed that TIP60 can interact with its cellular partner only in its phase-separated state, we wanted to investigate the physiological significance of TIP60’s phase separation in regulating TIP60-mediated cellular functions. Considering the importance of lysine 187 autoacetylation in regulating TIP60’s nuclear import and phase separation, we wanted to identify naturally occurring mutations in TIP60’s lysine 187 or adjacent amino acids and using cBioPortal (a cancer genomics portal)^35,36^, we identified two such naturally occurring mutations in the arginine 188 residue (R188P and R188Q) located adjacent to lysine 187 in renal clear cell carcinoma and colon adenocarcinoma, respectively (**Figure 6A**). In addition, we found two more mutations, including R177H (in cervical squamous cell carcinoma, head and neck squamous cell carcinoma and uterine endometrioid carcinoma) and R178H (in colorectal adenocarcinoma and melanoma) located in the nearby regions of lysine 187, within the IDR of TIP60. To investigate the effect of these mutations on TIP60’s localization and cellular functions, we first generated clones of TIP60 harbouring these mutations (R177H, R188H and R188P) in RFP-TIP60 (Wild-type) ORF background (**Figure S4C**). Western blot analysis was performed to confirm the expression of these mutant proteins (**Figure 6B**). To visualize the subcellular localization of these newly generated constructs, we also performed live cell imaging experiments with Cos-1 cells transfected separately with RFP-TIP60 (Wild-type), RFP-TIP60 (R177H), RFP-TIP60 (R178H) or RFP-TIP60 (R188P) plasmids. Interestingly, we found that out of all these mutants, TIP60 (R188P) mutant behaved similarly to the TIP60 (K187R) mutant and failed to enter the nucleus and remained in the cytosolic compartment (**Figure 6C**). However, under similar conditions two other mutants, RFP-TIP60 (R177H) and RFP-TIP60 (R178H), were found to form punctate nuclear foci similar to the Wild-type TIP60 protein.

**Figure 6.**
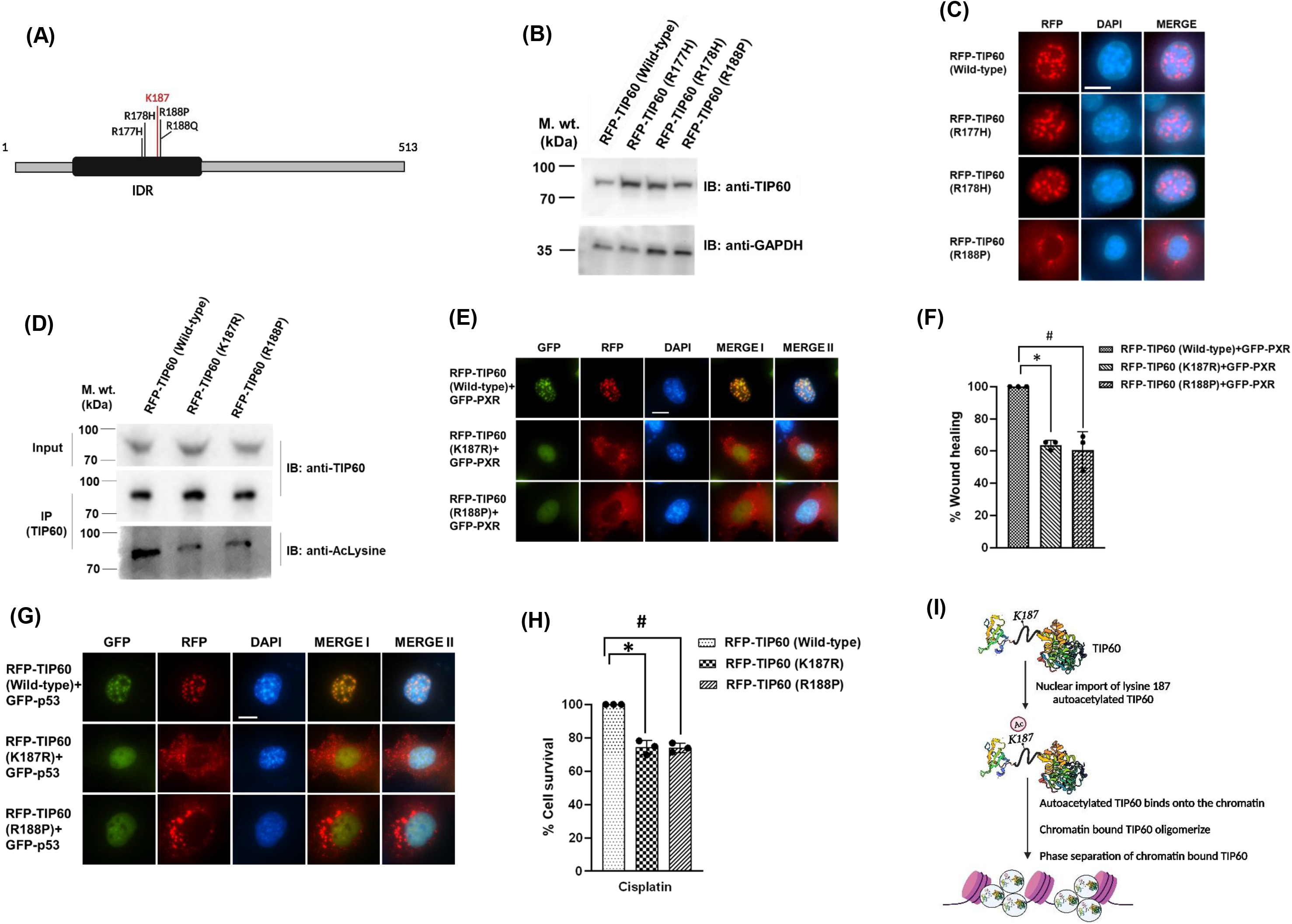
Analysis of cancer-associated mutations identified in TIP60’S IDR reveals importance of TIP60’s phase separation for its functions. **(A)** Schematic diagram showing the sites of cancer-associated mutations identified in TIP60’s IDR (black colour), from the cBioPortal database. Diagram is created with BioRender.com. **(B)** Western blot image showing the expression of cancer-associated IDR mutants. Cell lysates of transfected Cos-1 cells with indicated plasmids were resolved on SDS-PAGE followed by Western blotting with anti-TIP60 and anti-GAPDH antibody. **(C)** TIP60-IDR mutant (R188P) shows localization pattern similar to TIP60 (K187R) mutant. Live cell imaging was performed for Cos-1 cells transiently transfected with indicated plasmids to visualize the localization of fluorescent-tagged proteins, as indicated in red colour. DAPI is used as a nuclear stain (blue colour) and scale bar represents 10 µm. **(D**) TIP60-IDR mutant (R188P) shows diminished autoacetylation activity. RFP-TIP60 (Wild-type), RFP-TIP60 (K187R) or RFP-TIP60 (R188P) plasmids were transfected in Cos-1 cells followed by immunoprecipitation of TIP60 proteins using anti-TIP60 antibody. To detect the autoacetylation status, Western blot analysis was performed with anti-TIP60 or anti-acetylated lysine antibody. **(E)** TIP60-IDR mutant (K187R and R188P) fails to induce intranuclear reorganization of its interacting partner, PXR. Cos-1 cells were cotransfected with GFP-PXR and RFP-TIP60 (Wild-type) or RFP-TIP60 (K187R) plasmids followed by live cell imaging. Blue colour represents nucleus stained with DAPI. Green and red signals indicate the expression of GFP-tagged and RFP-tagged proteins. Merge I shows combined signals for TIP60 and PXR whereas merge II depicts signals for TIP60, PXR and DAPI. Scale bar shows 10 µm. **(F)** Wound healing capacity of TIP60-PXR complex was adversely compromised due to K187R or R188P mutations. HepG2 cells were transfected with the indicated plasmids followed by *in vitro* wound healing assay. Scratch was generated in transfected HepG2 cells and filling of wound gap was analyzed after 48 hours of wound generation. Graph generated using GraphPad Prism 8 software, represents the average value of three independent experimental replicates with S.D. p value for RFP-TIP60 (Wild-type)+GFP-PXR vs RFP-TIP60 (K187R)+GFP-PXR is 0.0021 (*), while p value for RFP-TIP60 (Wild-type)+GFP-PXR vs RFP-TIP60 (R188P)+GFP-PXR is 0.0272 (#). **(G)** TIP60-IDR mutants (K187R and R188P) fails to induce intranuclear reorganization of its interacting partner, p53. Cos-1 cells were transfected with indicated plasmids similar to (E) and live cell imaging was performed. Nucleus (blue colour) was stained with DAPI. Green and red colour signal indicate the expression of GFP-tagged and RFP-tagged proteins. Merge I shows combined signals for TIP60 and p53 whereas merge II depicts signals for TIP60, p53 and DAPI. Scale bar is equivalent to 10 µm. **(H)** DNA damage repair ability of TIP60 was adversely compromised due to K187R or R188P mutations. Huh-7 cells were transfected with the mentioned plasmids followed by cisplatin treatment for 6 hours. Cell viability were measured by counting the number of viable cells 48 hours post cisplatin withdrawal. The graph represents the average percentage of viable cells post-cisplatin treatment from three independent biological replicates with SD. p value for RFP-TIP60 (Wild-type) vs RFP-TIP60 (K187R) is 0.0091 (*), while p value for RFP-TIP60 (Wild-type) vs RFP-TIP60 (R188P) is 0.0037 (#). Graph was generated using GraphPad Prism 8 software. **(I)** Diagram showing the model of TIP60’s phase separation mechanism. TIP60 autoacetylate itself at lysine 187 residue which promotes its nuclear import. Inside the nucleus, autoacetylated TIP60 binds to the chromatin and oligomerize triggering its phase separation. Diagram is created with BioRender.com.

Since R188P mutation was located adjacent to lysine 187 (whose autoacetylation is critical for the phase separation of TIP60) we wanted to examine the impact of the R188P mutation on the autoacetylation status of the TIP60 protein. For this, Cos-1 cells were transfected with RFP-TIP60 (Wild-type) or RFP-TIP60 (K187R) or RFP-TIP60 (R188P) clones and TIP60 proteins were immunoprecipitated using anti-TIP60 antibody. Western blot analysis of these immunoprecipitated samples with anti-acetylated lysine antibody showed significant decrease in autoacetylation status of TIP60 (R188P) protein, similar to TIP60 (K187R) protein (**Figure 6D**). Together these results suggested that cancer-associated TIP60 mutations at arginine 188 residue resemble the features of the lysine 187 mutant and thus may affect the phase-separation of TIP60 in the respective cancers.

After determining the role of lysine 187 (autoacetylation site located in TIP60 IDR) and arginine 188 (a naturally occurring cancer-associated mutation identified in TIP60) residues in TIP60’s phase separation, we were now inclined to study the impact of these residues on TIP60’s cellular functions such as wound healing and DNA damage repair, involving its partner protein PXR and p53 respectively. Since TIP60 is known to interact with PXR and form colocalized nuclear foci during wound healing, we wanted to investigate the effect of these mutations on the formation of nuclear foci of TIP60 with PXR, followed by wound healing assay. First, we performed live cell imaging experiments with Cos-1 cells transfected with RFP-TIP60 (Wild-type) or RFP-TIP60 (K187R) or RFP-TIP60 (R188P) along with GFP-PXR plasmid and interestingly, found that both the mutants of TIP60 failed to form nuclear foci with PXR and remained in cytoplasm (**Figure 6E**). Subsequently, we performed wound healing assay using HepG2 cells transfected with indicated plasmids and a uniform scratch was generated in monolayer of transfected cells. Wound gap closure was monitored till 48 hours and graph was plotted for the obtained values. The results showed that the ability to migrate and fill the wound gap was significantly inhibited in cells overexpressing TIP60 (K187R) or TIP60 (R188P) mutants (**Figure 6F**).

Similarly, to investigate the effect of these mutations on TIP60 and p53-mediated cell survival during DNA damage condition, first we examined the effect of these mutations on the formation of nuclear foci of TIP60 with p53. For this, we performed live cell imaging experiment using Cos-1 cells transfected with GFP-p53 and RFP-TIP60 (Wild-type) or RFP-TIP60 (K187R) or RFP-TIP60 (R188P) plasmids. Interestingly, unlike Wild-type TIP60, both the mutants of TIP60 failed to enter the nucleus and did not form colocalized foci with p53 (**Figure 6G**). We next determined the effect of these mutations on TIP60-mediated DNA damage repair by performing cell survival assay using Huh-7 cells transfected with RFP-TIP60 (Wild-type) or RFP-TIP60 (K187R) or RFP-TIP60 (R188P) plasmids and subsequently treated with cisplatin (a DNA damaging agent), followed by counting of viable cells 48 hours post-treatment. The results showed significant reduction in the survival rate of cells expressing RFP-TIP60 (K187R) or RFP-TIP60 (R188P) mutants in comparison to cells expressing RFP-TIP60 (Wild-type), during DNA damage condition **(Figure 6H).** Together these findings suggest that phase separation of TIP60 is critical for interacting with its partner proteins and exerting the assigned cellular functions.

## DISCUSSION

To respond appropriately to the continual changes occurring in the cellular microenvironment, cells accordingly generate appropriate transcriptome mediated by the tightly coordinated and collaborative action of different transcription factors, histone modifying enzymes, chromatin remodellers and other co-regulatory factors at precise genomic targets^37–39^. Eukaryotic chromatin has an intricate structure which is dynamically organized to allow the expression of genes in a cell-type specific manner. Among various gene expression regulatory mechanisms, flexible compartmentalization of chromatin and assembly of chemically and structurally diverse biomacromolecules provide distinct intranuclear microenvironments to carry out and fine-tune distinct activities in the nucleus in spatio-temporal manner^5^. With the use of modern techniques such as chromosome conformation capture and high-resolution microscopy, we have been able to observe multiple levels of structuring of chromatin organization including extensive partitioning into active or inactive regions which may be further folded into self-interacting domains referred to as topologically associated domains (TADs)^40,41^. Studies have shown the association of TADs with several cis-regulatory elements (composed of promoters, enhancers and other genomic elements that control gene expression) forming chromatin hubs or condensates, important for regulating gene expression in a cell-type specific manner^42^. Depending on the type of interacting biomacromolecules, chromatin hubs can be of several types, such as chromatin-to-chromatin hubs, chromatin-to-large macromolecules, and chromatin-to-nuclear bodies^43^. For formation of these high order intranuclear membraneless spatial compartments, phase separation has emerged as a prominent phenomenon that allow low affinity multivalent interactions between different macromolecules within these chromatin hubs, to generate highly specialized environments optimized for specific biochemical processes^43,44^. It is believed that chromatin hub contains many transcription factors, chromatin remodellers and epigenetic regulators which help to attract and concentrate additional regulatory factors to rapidly regulate multiple gene expression^42^.

Interestingly, several studies have shown that transcription factors and coactivators like OCT4, GCN4, ER, Mediator, RNA polymerase II and MED1 makes phase separated condensates which enhances activation of their targeted genes^17,45,46^. Studies are now also showing the role of post-translational modifications of histones by epigenetic regulators in formation of these spatially and functionally distinct chromatin compartments^18,47^. Indeed, a recent report by Gibson et al showed that p300-mediated acetylation of histones disrupts chromatin phase separation while BRD4 (a multi bromodomain protein) was shown to promote LLPS of acetylated chromatin^18^. These findings exhibit the importance of histone modifiers/reader proteins and their role in segregating chromatin in phase-separated sub-compartments. In the present work, we have shown that TIP60, a chromatin modifier protein with crucial function in maintaining genomic integrity is capable of forming liquid-like condensates via liquid-liquid phase separation (LLPS) (**Figure 1**). We also found presence of an IDR in TIP60 that facilitates its phase separation in the nucleus, similar to many intrinsically disordered proteins (IDPs). Our observation that disruption of TIP60’s nuclear puncta by 1, 6-Hexanediol treatment does not hamper TIP60’s chromatin binding demonstrate that TIP60’s phase separation occur only after TIP60 is bound to the chromatin. TIP60 is known to catalyze the acetylation of its substrates including core histones through its HAT domain and also possesses a chromodomain that facilitates its binding to chromatin. Importantly, we also found in our study that, in addition to the indispensable requirement of IDR, TIP60’s chromatin binding ability as well as lysine acetyl transferase activity are highly essential for IDR-mediated TIP60 phase separation under *in vivo* condition. For example, we found that the chromatin binding ability of the TIP60 (HAT mutant) (with impaired catalytic activity) is significantly reduced which may be a major reason for the failure of TIP60 HAT mutant to form nuclear puncta^31^, which now we know are formed due to its phase separation. Similarly, we have recently shown that chromodomain mutant of TIP60 that severely loses its chromatin binding ability fails to form nuclear puncta^25^. These results clearly show us that the interplay of the different domains of TIP60 contributes to TIP60’s phase separation and TIP60’s chromatin binding is a pre-requisite step for its phase separation.

The role of PTMs in influencing protein stability, localization and interactions with partner proteins is not new^48^, but several studies now show that PTMs can also affect protein solubility and valency, which in turn can affect the phase separation of proteins^49,50^. PTMs can have diverse effects on IDP LLPS property, and can shift the equilibrium between the disordered and folded state potentially driven by stabilization/destabilization of local or global conformational changes, net charge of the protein, by altering their ability to interact with other proteins or by shifting the equilibrium between the state of dispersed monomeric and phase-separated IDPs^50,51^. For example, phosphorylation of the microtubule binding protein tau increases its propensity for phase separation while acetylation has been seen to disrupt its LLPS droplets^52^. Similarly, phosphorylation of FMRP enhance its phase separation while methylation, in contrast, attenuates its propensity for LLPS^53^. DDX3X, a component of stress granules (SGs) can also undergo LLPS and its acetylation at lysine K118 residue severely impairs its phase separation, whereas deacetylation by HDAC6 favours its liquid droplet formation, showing evidence of acetylation-deacetylation in modulating protein’s LLPS dynamics^54^. Since TIP60’s catalytic activity is also required for its autoacetylation, our analysis of the conserved autoacetylation sites of TIP60 revealed lysine 187 located within its IDR to be critical for TIP60’s phase separation. We found that lysine 187 is important for TIP60’s translocation into the nucleus and its oligomerization. A previous report had predicted that TIP60 contains a putative nuclear localization signal (NLS) and their anticipated NLS also included lysine 187 site^55^. However, the TIP60 mutant used in that study included three lysine mutations (located within putative NLS region) and in our analysis we did not found any impact of lysine 189 mutation on TIP60’s nuclear import and phase separation present within that predicted NLS region (Figure 2E). Thus we can speculate that the autoacetylation of lysine 187 may induce local structural modifications in its IDR which is critical for translocating TIP60 inside the nucleus. A previous example of similar kind shows, acetylation of lysine within the NLS region of TyrRS by PCAF promote its nuclear localization^56^. We also observed that catalytically comprised HAT mutant of TIP60 which do not form oligomers but can translocate inside the nucleus suggest that autoacetylation of TIP60 on lysine 187 helps to translocate monomeric TIP60 from cytosol into the nucleus where it then binds to the chromatin and oligomerize leading to its phase separation. Similar to the HAT mutant of TIP60 which could enter the nucleus but shows compromised chromatin binding ability, our fractionation data showed that K187R mutant also exhibited compromised chromatin binding ability compared to Wild type TIP60 (**Figure 2I**) indicating that TIP60 may contain additional post-translational modifications which may stabilize its chromatin binding.

Although our understanding of the functional diversity of phase-separated condensates and their physiological significance is insufficient at this stage, given the involvement of IDRs in maintaining cellular complexity and providing functional versatility for biochemical reactions, it is not surprising to see their involvement in development of many human diseases^57–59^. Aggregates of tau, β-amyloid and α-synuclein associated with various neurodegenerative diseases such as Alzheimer’s and Parkinson’s are actually found to be formed due to mutations in the IDRs of these proteins that increase the propensity to aggregate these IDPs^57^. Another example of a neurodegenerative disease associated with IDP is ALS (amyotrophic lateral sclerosis), which is caused by the formation of pathological agglutination of FUS, an RNA binding protein. Fus contains a prion-like domain that can form aggregates of the FUS protein. Patel A et al and colleagues showed in their study that patient-derived mutations in the FUS prion-like domain over time can cause abnormal phase transition of these protein, converting them from a liquid droplet to a fibrous state^60^. Several lines of evidence now exist linking aberrant phase separation to tumorigenesis^58,61^. Overexpression of SRSF1 (a component protein of NS) protein in breast cancer has been shown to be able to transform mammary epithelial cells and promote mammary gland tumorigenesis in mice^62^. Similarly, p53 which plays an important role in maintaining genomic integrity has been shown to form amyloid in human, mouse and rat cancer tissues^55^. In addition, amyloid formation promoted by cancer-associated mutations have also been found to impair the normal function of p53 in the cell^63^. It is now well established that transcription factors and coactivators can form phase-separated condensates at super enhancers in a cell-specific manner to regulate the gene expression and any type of interference with their phase-separation can lead to hyper transactivation of oncogenes^58,64^.

TIP60 has been shown to function primarily as a tumor suppressor in human cancers^65^. Mono-allelic loss of TIP60 has been shown to deregulate its expression level in several human cancers, rendering TIP60 incapable of exerting its tumor suppressor functions^65^. However, how various cancer-associated mutations identified in TIP60 impact or deregulate TIP60’s activity or its intracellular dynamics is not known. To investigate the effect of cancer-driven mutations on the phase separation and functions of TIP60, we specifically studied cancer-associated mutations located within the IDR of TIP60, where we found that arginine at position 188 got mutated to proline or glutamine, in two different cancer types (**Figure 6A**). From live cell imaging results, we found that the R188 mutant failed to enter the nucleus in the same way as the lysine 187 mutant and also affected TIP60’s ability to get autoacetylated. We also found additional similarities in their behaviour, as both the lysine 187 mutant and the cancer-associated R188 mutant neither formed nuclear puncta nor interacted with their interacting partners, such as PXR or p53 (**Figure 6E and 6G**). Moreover, these mutants behaved as functionally defunct TIP60 as they failed to promote TIP60-mediated wound healing or its DNA damage repair functions (**Figure 6F and 6H**). Our results clearly implicate inability of TIP60 to phase separate due to these mutations. These results are in sync with recently published study where authors have shown that p53 associates with NS and subset of its targeted gene’s including *p21* with NS, to enhance their transactivation^66^. It is well established that during DNA damage TIP60 regulates p53 to modulate cell equilibrium between cell cycle arrest or apoptosis by regulating the expression of *p21* or *PUMA*^67,68^. Also, in our recent study we showed that chromodomain mutant of TIP60 that fail to form nuclear puncta and display diminished histone acetylation activity could not transactivate its targeted genes during wound generated conditions^25^. Study by Gallego *et al* showed that phase separation of histone modifying enzymes promotes ubiquitination of histone H2B on gene-body nucleosome thus modulating their expression^69^. These examples help to postulate that phase separated TIP60 interact with its partner proteins and can stimulate the acetylation of histones within the condensates thus helps in modulating the targeted genes expression.

In the present work, we demonstrate that TIP60 undergoes phase separation to form liquid condensates in the nucleus that is highly required to perform its functions (**Figure 6I**). We identified the presence of an IDR in TIP60 that regulates its phase separation, however, our findings also highlighted that in addition to the IDR, the cooperative efforts of TIP60’s chromodomain as well as its HAT domain also play an important role in this phase separation. Our study uncovered another important aspect of TIP60, which suggests that TIP60 can interact with its interacting partner only in its phase-separated form. Our study reaffirms two important facts about IDPs and the condensates formed from them, that post-translational modifications act as critical regulators of phase separation ensuring that proteins segregate only into the required compartments so as to prevent their unwanted interactions or activities. Since IDRs do not yield a defined conformation, which provide promiscuity to IDPs for their selection of interacting partners and thus may confer multifunctionality to these proteins.

## Acknowledgements

SD and HG thanks ICMR for senior research fellowship. We acknowledge SNIoE for providing resources and space. Authors thank Prof. Sanjeev Galande for suggestions and encouragement and are highly obliged to Dr. Bharti Jaiswal for critical reading of the manuscript and providing insightful suggestions. Graphical abstract was created with BioRender.com.

## Author contributions

SD-performed most of the experiments in the study, original draft writing, editing, investigation, prepared the illustrations; HG-performed protein purification for LLPS, *in vitro* oligomerization and cell survivability experiments; AG-Conceived, designed and supervised the study, funding acquisition, original draft writing, review, and editing. AG, SD and HG participated in analyzing the data. All authors approved the final version of the manuscript.

## Material and Methods

### Cell Culture

Mammalian cell lines Cos-1, HepG2 and Huh-7 (procured from Cell Repository, NCCS, Pune, India) were cultured in Dulbecco’s Modified Eagle Medium (Gibco, USA; Hyclone, USA) supplemented with 0.5% pen/strep solution and 10% fetal bovine serum (Gibco, USA). Cells were cultured and maintained in a CO_2_ incubator under humid conditions, at 5% CO_2_ and 37□ temperature.

### Cloning

Autoacetylation mutants, ΔIDR mutant or cancer-associated mutants of TIP60 used in the study were generated using overlapping PCR method. Mutation (point or deletion) carrying amplicons were amplified using KpnI forward primer and mutation-specific reverse primer or mutation-specific forward primer and BamHI reverse primer. Amplified ORFs were then used as templates in a PCR reaction to amplify a full-length mutant amplicon followed by their purification using Wizard SV Gel and PCR Clean-Up kit (Promega, USA). These ORFs were digested with KpnI and BamHI enzymes and ligated into similarly digested pDsRed vector. *E. coli* DH5α cells were transformed with ligated products and screened for positive clones. Similarly, for cloning in pET28a bacterial expression plasmid, mutated amplicons (point or deletion) were amplified using forward primer (with BamHI restriction enzyme site) and reverse primer (with EcoRI restriction enzyme site).

### Transient transfection and live cell imaging

Cells were seeded in culture dish at 60-70% confluency and plasmids were transfected using lipofectamine 2000 reagent, following manufacturer’s protocol (Invitrogen, USA). After 5 hours of transfection, medium was replaced with DMEM containing 10% FBS. In case of PXR plasmid transfection, media was replaced with DMEM containing 10% stripped FBS. Post 24 hours of medium change, cells were processed for further experiments. To perform live cell imaging experiments, DAPI (Invitrogen, USA) was added at 0.5 µg/ml concentration and imaging of fluorescent tagged proteins was performed post 24 hours of medium change using Nikon Ti Eclipse fluorescence microscope (Nikon, USA). For 1, 6-Hexanediol (Sigma-Aldrich, USA) treatment, cells were treated with 3.5% 1, 6-Hexanediol for 5 minutes at room temperature, followed by live cell imaging or Western blot analysis.

### Immunofluorescence assay

For performing immunofluorescence assay, Cos-1 cells were seeded on glass coverslips and allowed to attach properly for 12 hours at 37[in CO_2_ incubator. Attached cells were washed thrice with 1X PBS and fixed using 4% formaldehyde for 15 minutes at room temperature. Fixed cells were permeabilized using 0.1% Triton X-100 for 5 minutes at room temperature, followed by washing with 1X PBS three times and subsequently incubated with 1% bovine serum albumin as blocking agent for 20 minutes at room temperature. Cells were then washed thrice with 1X PBS and incubated in primary antibody for 12 hours at 4[. Incubated cells were then washed thrice with 1X PBST (1X PBS with 0.1% tween 20) and incubated with Alexa 594 (anti-mice; Life Technologies, USA) or Alexa 488 (anti-rabbit; Life Technologies, USA) secondary antibodies at room temperature for 1 hour. Cells were again washed thrice with 1X PBST, air-dried and mounted with DAPI antifade mountant. Processed slides were then visualized using Zeiss LSM 800 confocal system (Zeiss, Germany).

### Immunoprecipitation assay

To perform immunoprecipitation assay, protocol was followed as mentioned elsewhere^25^. In brief, transiently transfected Cos-1 cells were lysed in the lysis buffer (20 mM Tris pH- 8.0, 2 mM EDTA, 150 mM NaCl, 0.5% Triton X 100, 0.1% SDS, and 1X protease inhibitor cocktail) for 1 hour at 4°C, followed by sonication at 30% amplitude for 2 cycles. The lysate was then centrifuged at 14,000 g for 10 minutes and 10% of total lysate was collected as input into the fresh vial. To prevent non-specific protein binding to the beads, lysate was precleared by incubating it with equilibrated protein-A Sepharose beads for 1 hour at 4°C. Precleared lysate was then mixed with anti-TIP60 antibody and equilibrated protein-A Sepharose beads and allowed to incubate overnight at slow speed. Beads were then centrifuged at 300 g for 5 minutes and washed twice in lysis buffer. The bead-bound immunoprecipitated proteins were eluted by adding 2X Laemmli buffer and proteins were resolved in SDS-PAGE followed by Western blotting analysis.

### Recombinant protein purification and pull down assay

For expression and purification of recombinant proteins from bacteria, *E. coli* BL21 DE3 cells were transformed with clones generated in pET28a or pGEX6p2 vectors. Recombinant TIP60 protein was expressed and purified using the protocol as described previously elsewhere ^25^. Briefly, transformed cells were cultured in Luria-Bertani broth at 37[, until O.D. reached 0.6. The culture was then treated with 0.5 mM Isopropyl-β-D-1-thiogalactopyranoside (IPTG) and protein expression was induced for 16 hours at 16[. Induced culture was pelleted at 3,000 g for 10 minutes and cells were lysed in lysis buffer (1X PBS, 2 mM EDTA, 5 mM DTT, 0.5 mM PMSF, 0.1% Triton X-100, 10% glycerol and 100 μg lysozyme) followed by sonication at 30% amplitude for 3 cycles. Sonicated samples were further lysed for 1 hour at 4[followed by centrifugation of lysate for 30 minutes at 14,000 g and equilibrated Ni-NTA beads were added to the clear supernatant along with 10 mM imidazole and incubated at slow speed for 1 hour. Protein-bound beads were centrifuged at 300 g for 5 minutes and washed twice with the wash buffer (1X PBS, 0.1 mM PMSF, 20 mM imidazole). Proteins were then eluted in elution buffer (50 mM Tris-pH 8.0, 10% glycerol, 150 mM NaCl, 500 mM imidazole, 1 mM PMSF) followed by dialysis in HAT buffer (50 mM Tris-pH 8.0, 10% glycerol, 0.1 mM EDTA, 1 mM DTT, 20 μM PMSF) at 4□.

To perform GST pull-down assay, *E. coli* BL21 DE3 cells were cotransformed with pGEX6p2-TIP60 (Wild-type) and pET28a-TIP60 (Wild-type) or pET28a-TIP60 (K187R) plasmids. Transformed cells were cultured under selection pressure of kanamycin and ampicillin and expression of recombinant proteins were induced using 0.5 mM IPTG at 0.6 O.D. Cells were lysed in lysis buffer (1X PBS, 2 mM EDTA, 5 mM DTT, 0.5 mM PMSF, 0.1% Triton X-100, 10% glycerol, 100 μg lysozyme and 10 µM acetyl-CoA) followed by sonication at 30% amplitude for 3 cycles. Sonicated samples were lysed in lysis buffer for 1 hour at 4[followed by centrifugation at 14,000 g for 30 minutes. Equilibrated glutathione S-transferase beads were added to the cleared supernatant and incubated at slow speed for 1 hour to facilitate binding of GST tagged proteins under native conditions. The protein bound GST beads were washed thrice in wash buffer (1X PBS, 0.1mM PMSF). Bound proteins were denatured in 2X Laemmli buffer and resolved in SDS-PAGE followed by Western blotting analysis.

### Western blot analysis

Protein samples prepared from cell lysates or purified recombinant proteins, were added with 2X Laemmli buffer and were resolved on SDS-PAGE followed by their transfer on methanol charged Polyvinylidene fluoride (PVDF) membrane using semidry transfer system (Biorad, USA). Membrane was further treated using 5% skimmed milk in PBST for 1 hour and then probed with primary antibody and incubated overnight at 4[. Membrane was then washed thrice with 1X PBST followed by incubation with secondary antibody was for 1 hour at room temperature. Incubated membrane was then washed thrice with 1X PBST and developed using enhanced chemiluminescence solution (Biorad, USA) using Protein Simple (FluorchemM, USA) or G:Box (Syngene, USA) gel documentation system.

### Histone acetyltransferase and autoacetylation assay

Histone acetyl transferase and autoacetylation assays were performed as mentioned previously^25^. In brief, *in vitro* HAT assay reaction was setup in HAT buffer added with purified recombinant proteins [(His-TIP60 (Wild-type) or His-TIP60 (K187R)], 0.5 µg histone H4 peptides (Millipore, USA) and 100 µM acetyl-CoA. Reaction mix was incubated at 30[for 1 hour and reaction was terminated by adding 2X Laemmli buffer. Samples were then resolved in 15% SDS-PAGE followed by Western blot analysis using anti-histone H4 (CST, USA) and anti-acetylated histone H4 (CST, USA) specific antibodies. Similarly, *in vitro* autoacetylation assay was performed by adding 1 µg of purified recombinant proteins [(His-TIP60 (Wild-type) or His-TIP60 (K187R)], 100 µM acetyl-CoA into HAT buffer and incubating the reaction mixture at 30[for 1 hour. Reaction samples were then resolved in 10% SDS-PAGE followed by Western blot analysis using anti-acetylated lysine antibody (Sigma, USA).

### Subcellular fractionation assay

For obtaining subcellular fractions, we followed the protocol mentioned elsewhere ^25^. Transfected cells were trypsinized and washed once with complete DMEM and twice with 1 X PBS, followed by solubilization using solubilisation buffer (10 mM HEPES pH-7.4, 10 mM KCl, 0.05% NP-40, 0.2 mM MgCl_2_, 1% Triton X 100, 100 mM NaCl, 1X protease inhibitor cocktail) for 20 minutes at 4□. Cells were centrifuged at 1,300 g for 5 minutes at 4[. The supernatant was gently aspirated and aliquoted separately as soluble fraction (containing cytoplasm and nucleoplasm fractions). The pellet was further washed twice with solubilisation buffer and suspended in chromatin buffer (50 mM Tris pH-8.0, 400 mM NaCl, 10 mM EDTA, 0.5% SDS, 1X protease inhibitor cocktail) for 20 minutes at 4□, followed by sonication at 20% amplitude for 2 cycles. The sonicated samples were centrifuged at 1,700 g for 5 minutes and supernatant was collected as chromatin fraction. Further, Bradford reagent was used for quantifying both soluble or chromatin fractions and 10 µg/µl of protein sample of each fraction was denatured in 2X Laemmli buffer followed by Western blot analysis.

### *In vitro* LLPS formation, turbidity assay and oligomerization assay

For *in vitro* LLPS formation, recombinant proteins were purified in a similar manner as described previously with some modifications (lysis buffer contains 500 mM NaCl, wash buffer contain 30 mM imidazole and elution buffer contains 500 mM NaCl in addition to other constituents). To detect *in vitro* liquid-liquid phase separation, the indicated concentrations (1 µM, 5 µM, 10 µM, 15 µM) of purified recombinant proteins were incubated with PEG 8000 (at different percentages, (w/v)) followed by final dilution in phase separation buffer (50 mM Tris-pH 8.0, 10% glycerol, 500 mM imidazole, 1 mM PMSF). Samples were then visualized using Nikon Ti Eclipse fluorescent microscope (Nikon, USA) for differential interference contrast (DIC) microscopy. Turbidity assay was performed to evaluate phase separation of His-TIP60 (Wild-type) and His-TIP60 (ΔIDR) proteins. Different concentrations of purified proteins (1 µM, 5 µM, 10 µM, 15 µM) were incubated with 15% PEG 8000 followed by final dilution in phase separation buffer. The optical density of solution was measured at OD_600_ nm using PerkinElmer multimode plate reader (PerkinElmer, USA).

To perform *in vitro* oligomerization assay, recombinant His-tagged TIP60 (Wild-type), His-TIP60 (HAT mutant) or His-TIP60 (K187R) were crosslinked using glutaraldehyde as a crosslinking agent. Purified proteins were incubated with 0.00025% or 0.0005% glutaraldehyde solution (in 1X PBS) for 20 minutes at 37[. To stop the crosslinking reaction, 2X Laemmli buffer was added and samples were resolved in 8% SDS-PAGE, followed by Western blot analysis using anti-His antibody (Invitrogen, USA).

### Wound healing assay and cell survival assay

For performing wound healing assay, HepG2 cells were seeded at 80% confluency and were transfected with the respective plasmids. Post 24 hours of medium change, a scratch was generated using 10 µl pipette tip. The gap filling of the generated wound was measured and high-resolution images were captured at regular time intervals till 48 hour using Nikon Ti Eclipse fluorescence microscope (Nikon, USA). Different wound areas were measured using NIS elements analysis tool provided with the system. The average values measured were further used to calculate % wound healing using the formula mentioned below:

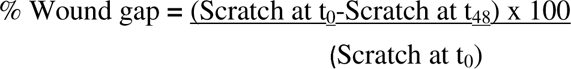

To examine the effect of cancer-associated mutations on TIP60’s DNA damage repair function, a cell survival assay was performed with Huh-7 cells, for the same 50,000 cells/ well were seeded in a 12-well plate, followed by transfection with RFP-TIP60 (Wild-type), RFP-TIP60 (K187R), and RFP-TIP60 (R188P) plasmids. Then, the cells were treated with cisplatin (5 µM concentration) for 6 hours, and subsequently, the medium was changed to DMEM containing 10% FBS. 48 hours post-treatment, viable cells were measured using a hemocytometer.

### Fluorescence recovery after photobleaching (FRAP)

To conduct a FRAP, Cos-1 cells were seeded in a 35 mm glass bottom dish and transfected with pDsRed-TIP60 plasmid, followed by time-lapse imaging of cells expressing RFP-TIP60 using a Nikon Ti2 Eclipse confocal microscope (Nikon, USA). RFP-TIP60 puncta’s were bleached with 100% laser power at 561 nm and recovery was recorded at 50 seconds per frame for 10 minutes. The intensities of bleached and post-bleach florescence signal recovery were normalised with respect to the maximum fluorescence intensity before bleaching (in %) for 20 independent puncta.

### Multiple sequence alignment and Statistical analysis

To examine whether lysine 187 residue is conserved in TIP60, sequences for TIP60 protein from various species were retrieved from protein database of NCBI and multiple sequence alignment analysis was performed using Clustal Omega tool.

Statistical significance between two samples was determined using unpaired t-test with Welch’s correction whereas for multiple samples, significance was calculated using one-way ANOVA with Bonferroni’s multiple comparison test. P value, p <0.05 was considered significant for the analysis. GraphPad Prism 8 software was used to create graphs displaying mean values (SD) for three independent experimental replicates.

