## Supplementary Information for "Autoacetylation-mediated phase separation of TIP60 is critical for its functions"

### Supplemental information

#### Figure legends

**Figure S1. (A)** The amino acid sequence of TIP60 was analyzed for the presence of intrinsically disordered region (IDR). IDR in TIP60 protein was predicted using dSCOPE webserver between 68 to 222 amino acid region. **(B)** IDR prediction was performed using VL3 predictor of PONDR. The prediction showed the region covering 79-205 amino acids as disordered. **(C)** Primary sequence of full-length TIP60 protein was analyzed for its droplet promoting ability. Region covering 64-219 amino acids in TIP60 was shown as droplet promoting regions (DPR).

**Figure S2. (A)** Differential interference contrast (DIC) images depicting phase-separated liquid droplets of His-TIP60 (Wild-type) at different protein concentrations (1 µM, 5 µM, 10 µM, 15 µM). Different concentrations of PEG 8000 (0%, 10% and 15%) were used as crowding agent in the experiment. Scale bar is 5 µm. **(B)** Cos-1 cells were transfected with RFP-TIP60 (Wild-type) or RFP-TIP60 (ΔIDR) and cells were harvested after 24 hours of medium change. After lysing the cells, protein samples were treated with 2X Laemmli buffer and resolved using SDS-PAGE. Western blot analysis was performed using anti-TIP60 and anti-GAPDH antibody. **(C)** Cos-1 cells were transfected with RFP-TIP60 (Wild-type) plasmid and after 24 hours of medium change, cells were treated with 3.5% 1, 6-Hexanediol for 5 minutes. The harvested cells were lysed and protein samples were resolved in SDS-PAGE followed by Western blotting using anti-TIP60 and anti-GAPDH antibody. **(D)** Multiple sequence alignment of TIP60 protein sequences for different species was performed using Clustal Omega tool. Green colour shows conserved lysine residue, corresponding to hTIP60 lysine 187, in vertebrates.

**Figure S3. (A)** Coomassie gel representing bacterially purified recombinant His-TIP60 (Wild-type), His-TIP60 (HAT mutant) and His-TIP60 (K187R) proteins. **(B)** Western blot showing expression of full length GFP-TIP60. Cos-1 cells were transiently transfected with generated GFP-TIP60 plasmid and untransfected cells were taken as control. Cells were lysed and protein samples were treated with 2X Laemmli buffer. Samples were resolved in SDS-PAGE followed by Western blotting analysis using anti-TIP60 and anti-GAPDH antibody.

**Figure S4. (A)** Cos-1 cells were transfected with RFP-TIP60 (Wild-type) or RFP-TIP60 (HAT mutant) and GFP-PXR plasmids, as indicated in the figure panel, followed by live cell imaging of fluorescent tagged proteins. Green and red signals indicate the expression of GFP tagged and RFP tagged proteins while blue colour shows nucleus stained with DAPI. Merge I represents the combined signals for GFP and RFP while merge II shows combined signals for GFP, RFP and DAPI. Scale bar is 10 µm. **(B)** PXR interact with wild-type TIP60 but not with its catalytically compromised mutant form. Cos-1 cells were transfected with indicated plasmids and cells were harvested after 24 hours of medium change. Harvested cells were lysed and immunoprecipitation assay was performed using PXR antibody and immunoprecipitated samples were resolved in SDS-PAGE. Western blot analysis was performed using anti-TIP60 and anti-PXR antibody. **(C)** Overlapping PCR for cancer-associated TIP60 mutations observed in IDR region. 1% agarose gel representing the amplification of PCR-I, PCR-II, and full-length PCR-III for TIP60's R177H, R178H and R188P mutants.

**Table 1: List of primers used in the study.**

| **S. No.** | **Primer** | **Sequence (5'-3')** |
|---|---|---|
| 1 | hTIP60-KpnI-Fw | GG**GGTACC**ATGGCGGAGGTGGGGGAG |
| 2 | hTIP60-BamHI-Rv | CG**GGATCC**TCACCACTTCCCCCTCTTGC |
| 3 | hTIP60-BamHI-Fw | CG**GGATCC**ATGGCGGAGGTGGGGGAG |
| 4 | hTIP60-EcoRI-Rv | CG**GAATTC**CCACTTCCCCCTCTTGCT |
| 5 | hTIP60-EcoRI-Fw | CG**GAATTC**ATGGCGGAGGTGGGGGAG |
| 6 | hTIP60-K76R-Fw | AAGAAAGAGGCCCGGACCCCCACT |
| 7 | hTIP60-K76R-Rv | CTTAGTGGGGGTCCGGGCCTCTTT |
| 8 | hTIP60-K80R-Fw | AAGACCCCCACTCGGAACGGACTT |
| 9 | hTIP60-K80R-Rv | AGGAAGTCCGTTCCGAGTGGGGGT |
| 10 | hTIP60-K148R-Fw | GAGAGAGAGGTGCGACGGAAGGTG |
| 11 | hTIP60-K148R-Rv | CTCCACCTTCCGTCGCACCTCTCT |
| 12 | hTIP60-K150R-Fw | GAGGTGAAACGGCGGGTGGAGGTG |
| 13 | hTIP60-K150R-Rv | AACCACCTCCACCCGCCGTTTCAC |
| 14 | hTIP60-K187R-Fw | CAGCCAGGACGGCGGCGAAAATCG |
| 15 | hTIP60-K187R-Rv | ATTCGATTTTCGCCGCCGTCCTGG |
| 16 | hTIP60-K189R-Fw | GGACGGAAGCGACGATCGAATTGT |
| 17 | hTIP60-K189R-Rv | CAAACAATTCGATCGTCGCTTCCG |

| | | |
|---|---|---|
| 18 | hTIP60-K327R-Fw | CTTGACCATAGGACACTGTACATAT |
| 19 | hTIP60-K327R-Rv | GTACAGTGTCCTATGGTCAAGGAA |
| 20 | hTIP60-K383R-Fw | CGGGGCTACGGCCGGCTGCTGATC |
| 21 | hTIP60-K383R-Rv | CTCGATCAGCAGCCGGCCGTAGCC |
| 22 | hTIP60-R177H-Fw | AATGGAGCCGCCCATAGGGCAGTG |
| 23 | hTIP60-R177H-Rv | TGCCACTGCCCTATGGGCGGCTCC |
| 24 | hTIP60-R178H-Fw | GGAGCCGCCCGTCATGCAGTGGCA |
| 25 | hTIP60-R178H-Rv | GGCTGCCACTGCATGACGGGCGGC |
| 26 | hTIP60-R188P-Fw | CCAGGACGGAAGCCAAAATCGAAT |
| 27 | hTIP60-R188P-Rv | ACAATTCGATTTTGGCTTCCGTCC |
| 28 | hTIP60-ΔIDR-Fw | ACCCCCACTAAGACCCGGATGAAG |
| 29 | hTIP60-ΔIDR-Rv | CTTCATCCGGGTCTTAGTGGG |

#### Supplementary Figure S1

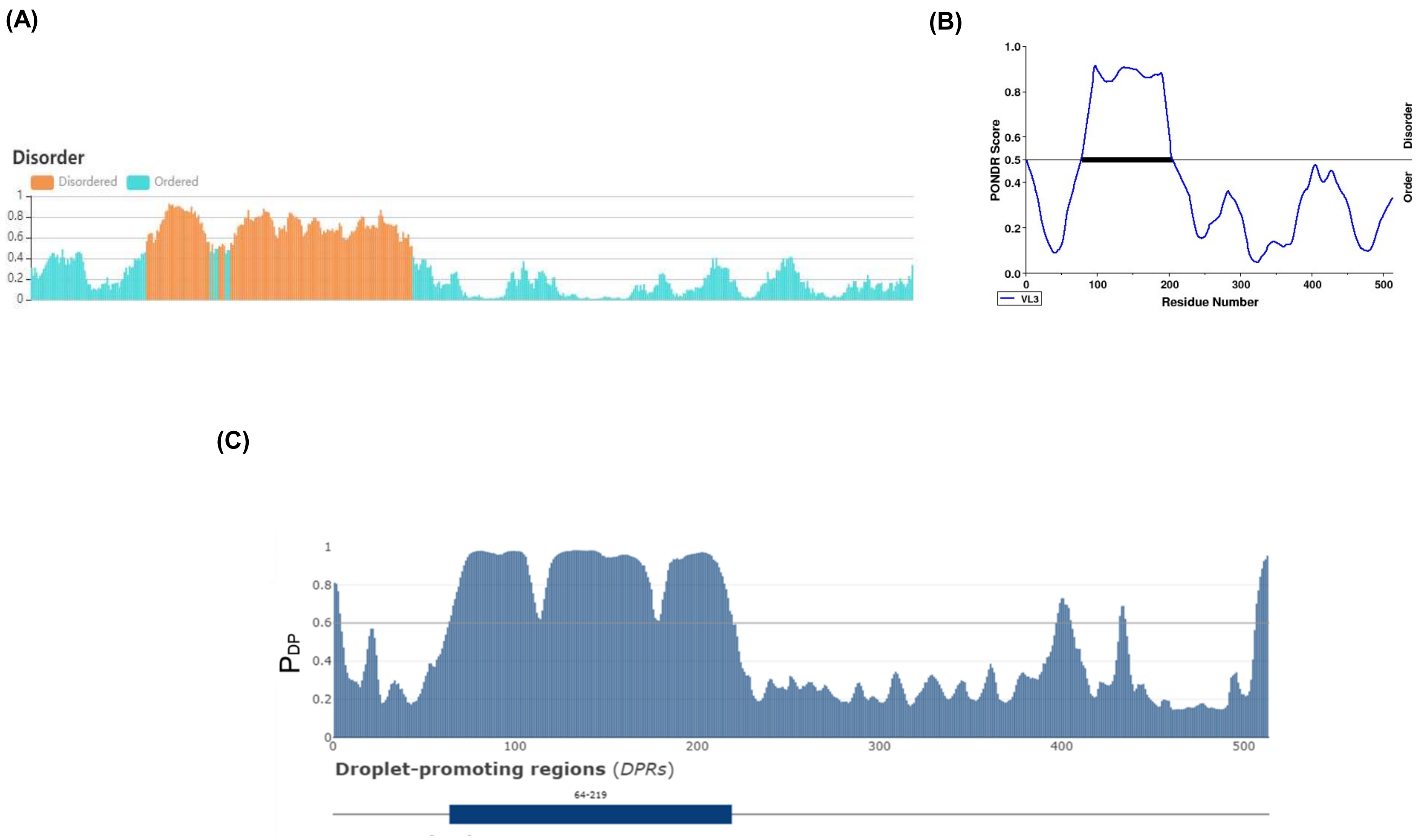

#### Supplementary Figure S2

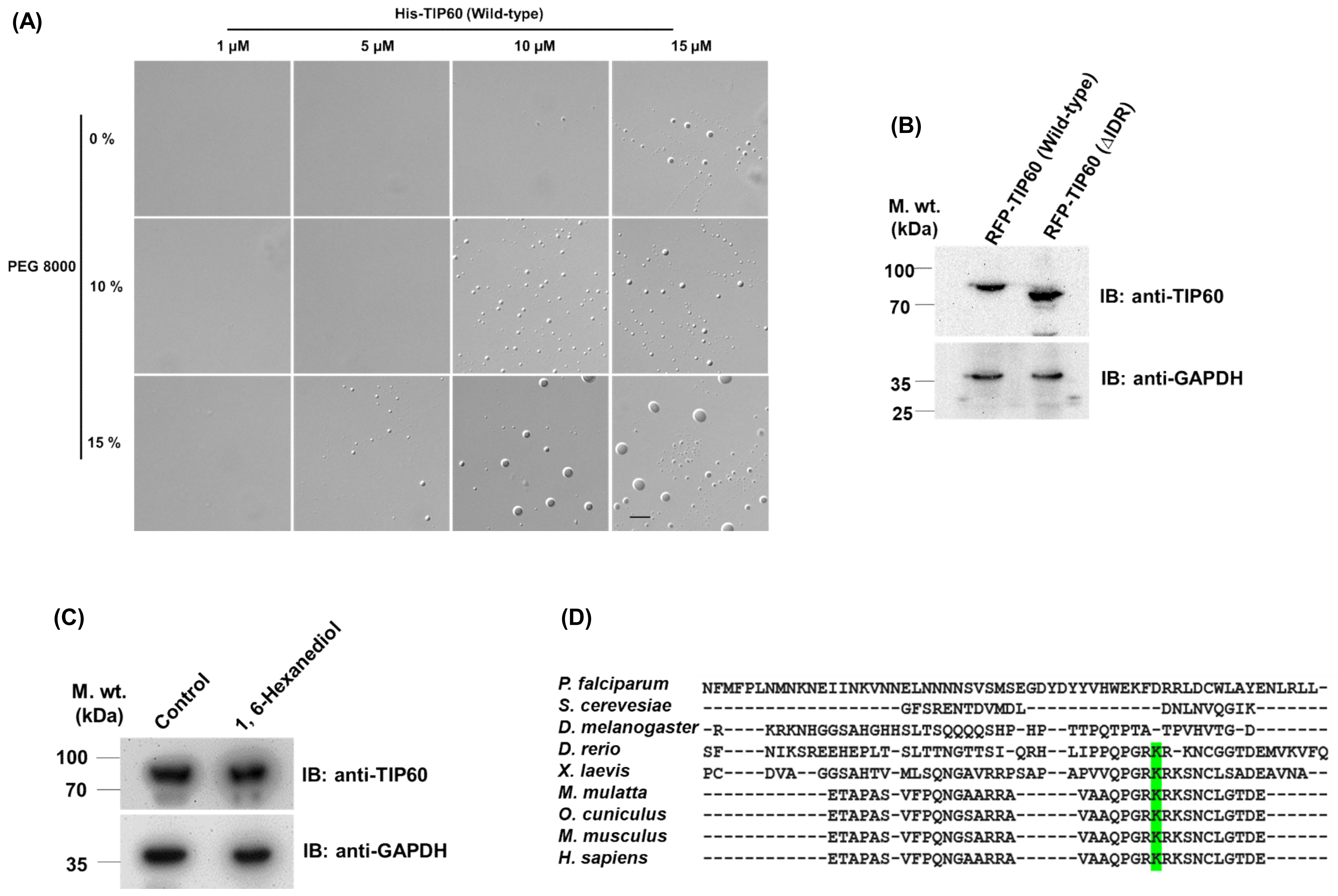

#### Supplementary Figure S3

(A)

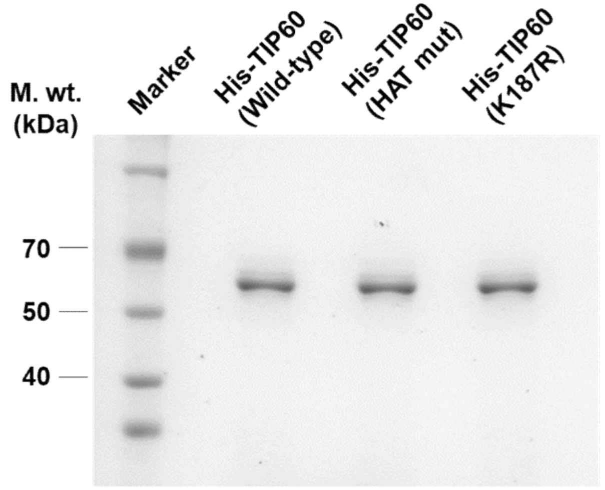


(B)

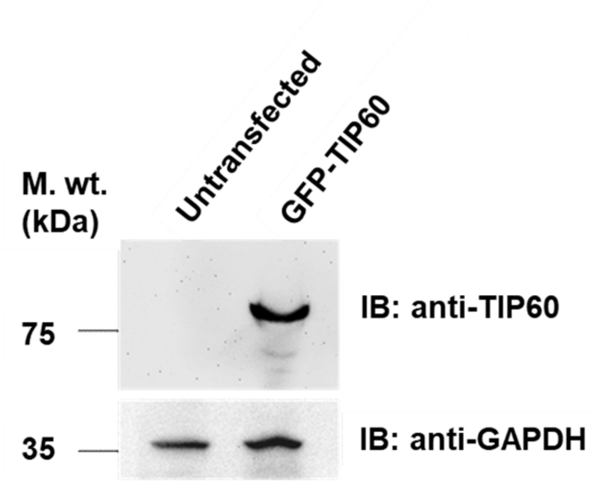

### Supplementary Figure S4

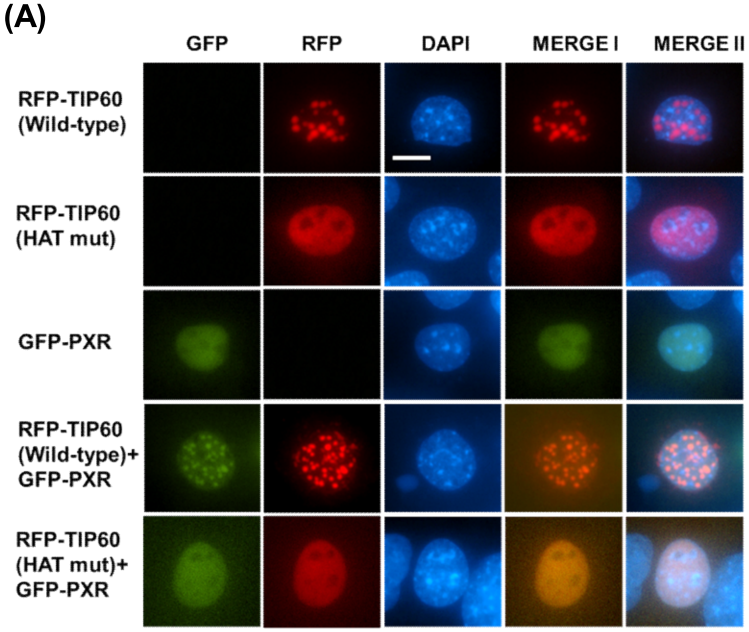


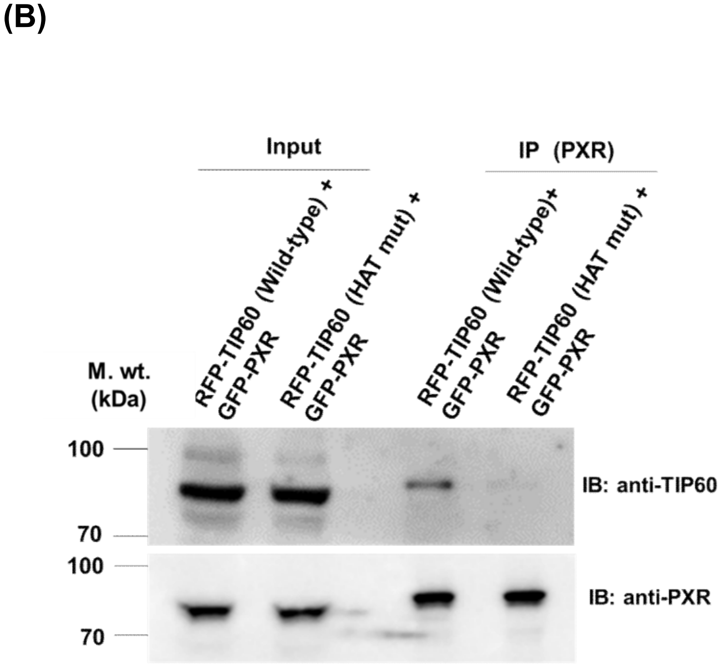


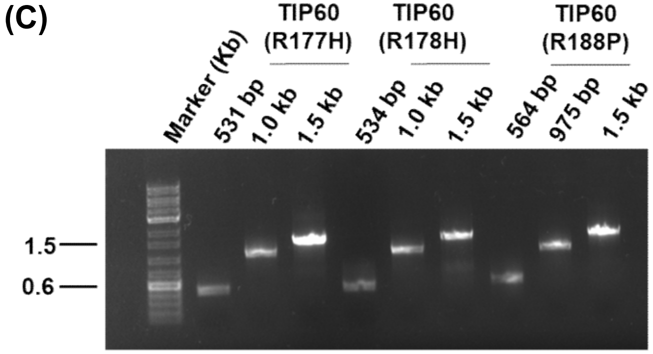
